# Orientation-Independent-DIC imaging reveals that a transient rise in depletion force contributes to mitotic chromosome condensation

**DOI:** 10.1101/2023.11.11.566679

**Authors:** Shiori Iida, Satoru Ide, Sachiko Tamura, Tomomi Tani, Tatsuhiko Goto, Michael Shribak, Kazuhiro Maeshima

## Abstract

Genomic information must be faithfully transmitted into two daughter cells during mitosis. To ensure the transmission process, interphase chromatin is further condensed into mitotic chromosomes. Although protein factors like condensins and topoisomerase IIα are involved in the assembly of mitotic chromosomes, the physical bases of the condensation process remain unclear. Depletion force/macromolecular crowding, an effective attractive force that arises between large structures in crowded environments around chromosomes, may contribute to the condensation process. To approach this issue, we investigated the “chromosome milieu” during mitosis of living human cells using orientation-independent-differential interference contrast (OI-DIC) module combined with a confocal laser scanning microscope, which is capable of precisely mapping optical path differences and estimating molecular densities. We found that the molecular density surrounding chromosomes increased with the progression from prometaphase to anaphase, concurring with chromosome condensation. However, the molecular density went down in telophase, when chromosome decondensation began. Changes in the molecular density around chromosomes by hypotonic or hypertonic treatment consistently altered the condensation levels of chromosomes. *In vitro*, native chromatin was converted into liquid droplets of chromatin in the presence of cations and a macromolecular crowder. Additional crowder made the chromatin droplets stiffer and more solid-like, with further condensation. These results suggest that a transient rise in depletion force, likely triggered by the relocation of macromolecules (proteins, RNAs and others) via nuclear envelope breakdown and also by a subsequent decrease in cell-volumes, contributes to mitotic chromosome condensation, shedding light on a new aspect of the condensation mechanism in living human cells.

**Significance Statement:** Mitotic chromosome condensation is an essential process to transmit replicated chromosomes into two daughter cells during cell division. To study the underlying physical principles of this process, we focused on depletion force/macromolecular crowding, which is a force that attracts large structures in crowded cell environments. Using newly developed special light microscopy, which can image the molecular density of cellular environments, we found that crowding around chromosomes increases during cell division. *In vitro*, higher concentrations of macromolecules condense chromatin and make it stiffer and more solid-like. Our results suggest that the rise in depletion force renders chromosomes more rigid, ensuring accurate chromosome transmission during cell division.

## Introduction

Negatively charged genomic DNA wraps around basic core histone proteins to form nucleosomes. The string of nucleosomes, together with other nonhistone proteins and RNAs, are somewhat irregularly organized in the cell as chromatin (1, 2). During interphase in higher eukaryotic cells, chromatin forms condensed domains as their functional units (3–5). With cell cycle progression, genomic DNA is duplicated by DNA replication and faithfully transmitted into the two daughter cells during mitosis (6, 7). To ensure the transmission process, interphase chromatin is further condensed into mitotic chromosomes. While several players involved in the condensation process, including condensins and topoisomerase IIα, have been identified and extensively investigated (8–13), physical bases of the condensation process remain unclear (14, 15).

In addition to these protein factors, two kinds of physical forces governed by the chromosome milieu may contribute to the condensation process. First, free divalent cations such as Mg^2+^ condense chromatin or chromosomes *in vitro* (16–23), or by computer modeling (24). These cations have long been considered important for local condensation (i.e., nucleosome-nucleosome interactions) because the nucleosomes have a net negative charge and are stretched like “beads on a string” by an electrostatic repulsion force in the absence of cations. Mg^2+^ counters this negative charge and repulsion force. Indeed, a transient increase in free Mg^2+^ is observed during mitosis, contributing to chromosome condensation (25). In addition to Mg^2+^, acetylation/deacetylation of histone H3 and H4 tails also regulates the electrostatic force between nucleosomes (15, 26).

The other possible physical force is the depletion force/macromolecular crowding effect (27–30). This force may also be critical because the cellular environment is highly crowded with macromolecules such as proteins, RNA, DNA, and others resulting in a molecular density >100 mg/mL. The principle of depletion force/macromolecular crowding effect is simple (Schematic in Fig.1A). Large cellular complexes, like chromatin, are bombarded from all sides by many soluble macromolecules in a crowded environment. When two large complexes come into contact, the macromolecules exert a force equivalent to their osmotic pressure on opposite sides of the two large complexes to keep them together (Schematic in Fig.1A) (27–29). This state is entropically favored because there are more accessible regions after their contact (Schematic in Fig.1A). Indeed, purified nuclei (31) and chromosomes (32) condensed *in vitro* with ∼10% (w/v) of long neutral polymers such as polyethylene glycol (PEG) or polyvinyl alcohol. Such high concentrations of a macromolecular “crowder” induced folding of long synthetic chromatin fibers *in vitro* (33). When ∼10% of bovine serum albumin (BSA; an anionic protein) was used as a crowder, condensates of the synthetic chromatin fibers were observed *in vitro* (34). However, it remains unclear whether the depletion force/macromolecular crowding effect is really involved in mitotic chromosome condensation in the cell, especially in higher eukaryotic cells.

**Figure 1.**
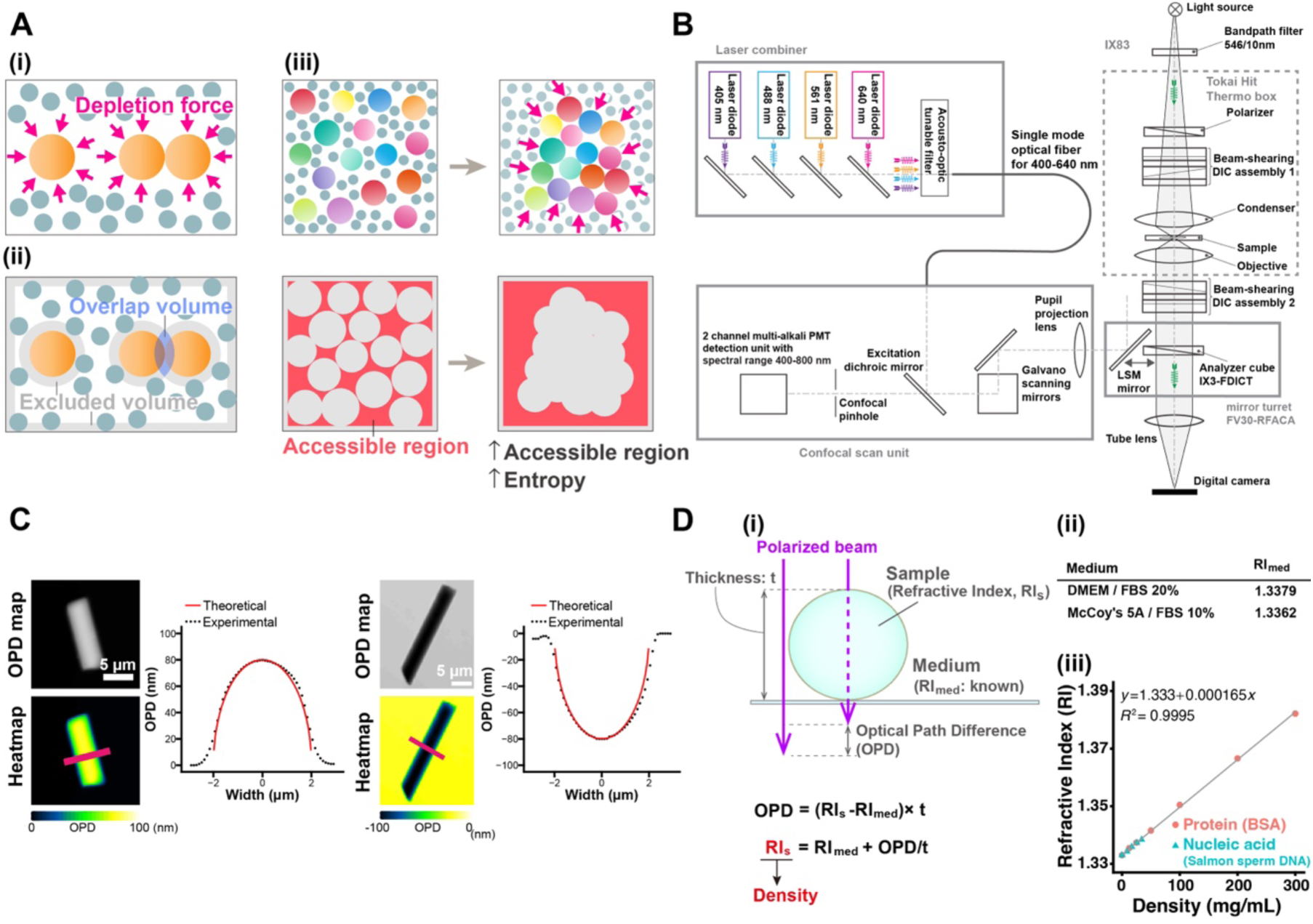
Schematics of macromolecular crowding effect/depletion force, OI-DIC microscopy, and density quantification. (**A**) (i) Many small spheres (blue-gray) representing soluble macromolecules bombard three large spheres (orange), representing chromatin, from all sides (arrows). When two large spheres come into contact (right), the small ones exert a force equivalent to their osmotic pressure on opposite sides of the two large ones to keep them together (macromolecular crowding effect/depletion force). (ii) The shaded regions in this alternative view show regions inaccessible to the centers of mass of the small spheres (excluded volumes). When two large spheres contact each other, their excluded volumes overlap and increase the volume accessible to the small spheres. Aggregation of the large spheres is then favored by an increase in the entropy of the system. (iii) Chromatin condensation by macromolecular crowding/depletion force. Chromatin domains/globules (colored spheres) are present in a square. The excluded volume is shown in gray. If chromatin domains associate with each other, the accessible region for soluble macromolecules (red) increases (excluded volume decreases). This state is then favored. **(B)** Optical schematic of confocal laser scanning microscope Olympus FV3000 equipped with OI-DIC module. Details of the microcopy system are described under SI Materials and Methods. **(C)** Validation of density imaging by OI-DIC microscopy using known glass rods and mineral oils. The RI of the glass rods was 1.56, and those of the oils were 1.54 (left) and 1.58 (right). Note that the theoretical and experimental values are almost the same, ensuring the accuracy of our RI quantification. Scale bar: 5 μm. **(D)** (i) A procedure for estimating the RI of sample (depicted as a sphere). Our OI-DIC microscopy can computationally quantify optical path differences (OPDs) at each spatial point. For details, see SI Materials and Methods. (ii) The RI of medium measured by the refractometer. (iii) The calibration curve of RI versus the density of standard solutions for protein or nucleic acid. RI = 1.333 + 1.65 × 10^-4^ × C (RI, the refractive index; C, the concentration of the proteins or nucleic acids [mg/mL]). For details, see SI Materials and Methods.

To observe the depletion force /macromolecular crowding effect on chromosome condensation, it is essential to measure molecular density in the chromosome milieu of living cells. While it is technically challenging, we gain insight on molecular density by using differential interference contrast (DIC) microscopy (35–38), a standard tool in cell biology research. DIC images are produced by the interference of two laterally displaced light beams passing through a sample (e.g., live cells), capturing information about the optical path length (OPL) in the sample to reveal otherwise invisible features (35). The difference in OPL (optical path difference, OPD) between the two beams contrasts the image, reflecting local differences in the refractive index (RI) within the sample. However, the contrast in DIC images depends on the direction of displacement between the two light beams and the sample, precluding quantitative measurement of the OPL. To overcome the limitations of DIC, Shribak et al. developed an orientation-independent-DIC (OI-DIC) microscopy method (39), which allows the directions of displacement for the two light beams to be rapidly switched by 90° without mechanically rotating the sample or the prisms, generating a quantitative OPD map (Fig. S1). Based on the OPD value, it is possible to estimate the density of intracellular regions in live cells. However, a previous type of OI-DIC with a wide-field epi-fluorescence microscope had difficulty in measuring precise thickness of target structures in the cell (40, 41).

In this study, we have combined a confocal laser scanning microscope (CLSM) with an OI-DIC module (Fig. 1B), which can provide high-resolution maps of OPD by OI-DIC and the thickness of target structures in the cell by CLSM. This new OI-DIC imaging can generate a 3D volume image of the RI and molecular density (dry mass) in living cells based on the calibration data (Fig. 1D(iii)). Using this new OI-DIC system, we quantified the absolute density of the molecules around chromosomes during mitosis of human HCT116 cells. We found that the molecular density surrounding mitotic chromosomes increased with the mitotic progression from prometaphase to anaphase, concurring with chromosome condensation. In telophase, the molecular density decreased as chromosomes began to decondense. Hypertonic treatment of the mitotic cells rapidly increased density and induced chromosome condensation, while hypotonic treatment had the opposite effects. *In vitro*, fiber-like condensates of native chromatin, induced by physiological concentrations of cations, were efficiently converted into chromatin liquid droplets with macromolecular crowders such as PEG, BSA, or Dextran. Additional amounts of crowders made the chromatin droplets stiffer and more solid-like structures. These results suggest that during mitosis a transient rise in depletion force by proteins, RNAs, and others contributes to mitotic chromosome condensation, shedding light on a new aspect of mitotic chromosome condensation in living human cells.

## Results

### Development of a new OI-DIC system combined with a confocal laser scanning microscope

To estimate the density of total molecules in the chromosome milieu, we developed a new OI-DIC module combined with a confocal laser scanning microscope (CLSM) (Fig. 1B). We calculated the dry mass density of the sample (Fig. 1D(i); for more details, see SI Materials and Methods) based on the OPD map obtained from OI-DIC imaging (e.g., Figs. 1C and S1), the precisely measured thickness (t) of the target structure by CLSM (Figs. 1D, S2C, S5A, and S5C), and the known refractive index (RI) of the surrounding medium. Next, we evaluated whether the new OI-DIC imaging and subsequent analysis could accurately estimate RIs by observing glass rods (diameter = 4 µm) in mineral oils with known refractive indices (RI = 1.54 and 1.58) and calculated the theoretical OPD (Fig. 1C). The theoretical and optically measured OPDs were almost identical (Fig. 1C), validating the accuracy of our new OI-DIC imaging for estimating the RI of samples with measured OPD.

### OI-DIC imaging of interphase and mitotic live human HCT116 cells

Using the procedure described above (Fig. 1D), we performed OI-DIC imaging of interphase and mitotic live human HCT116 cells stably expressing H2B-HaloTag and obtained their OPD maps (Fig. 2A). Riesz images are edge-enhanced OPD maps according to the inverse Riesz transform (42, 43) and are more visually informative with the higher contrast of small details (bottom images, Fig. 2A). Note that asynchronous live cells were used for this imaging to avoid any artifactual effects, such as accumulations and aggregations of proteins and RNAs, caused by the synchronization process. We clearly observed cytoplasmic organelles, nuclear envelopes, and presumably nucleoli in OPD maps of interphase HCT116 cells (Fig. 2A, lower left). In the case of mitotic cells, cytoplasmic organelles, and condensed chromosomes were notable in the Riesz images (Fig. 2A, bottom images). We set several ROIs in each cell to avoid apparent structures in the Riesz image, presumably endoplasmic reticulum, Golgi apparatus, and mitochondria (Fig. S3) (44). Cytoplasmic ROIs were set to exclude chromosomes in the entire Z-stacks (Fig. S3). To obtain the thickness measurement (t) of cytoplasm (Figs. 2B, S2A, S2C, and S5A) and chromatin (for nucleoplasm, see Figs. S2B and S2C; for chromosomes, Figs. S5B and S5C) by CLSM, cells were simultaneously labeled with Calcein AM and HaloTag ligand TMR.

**Figure 2.**
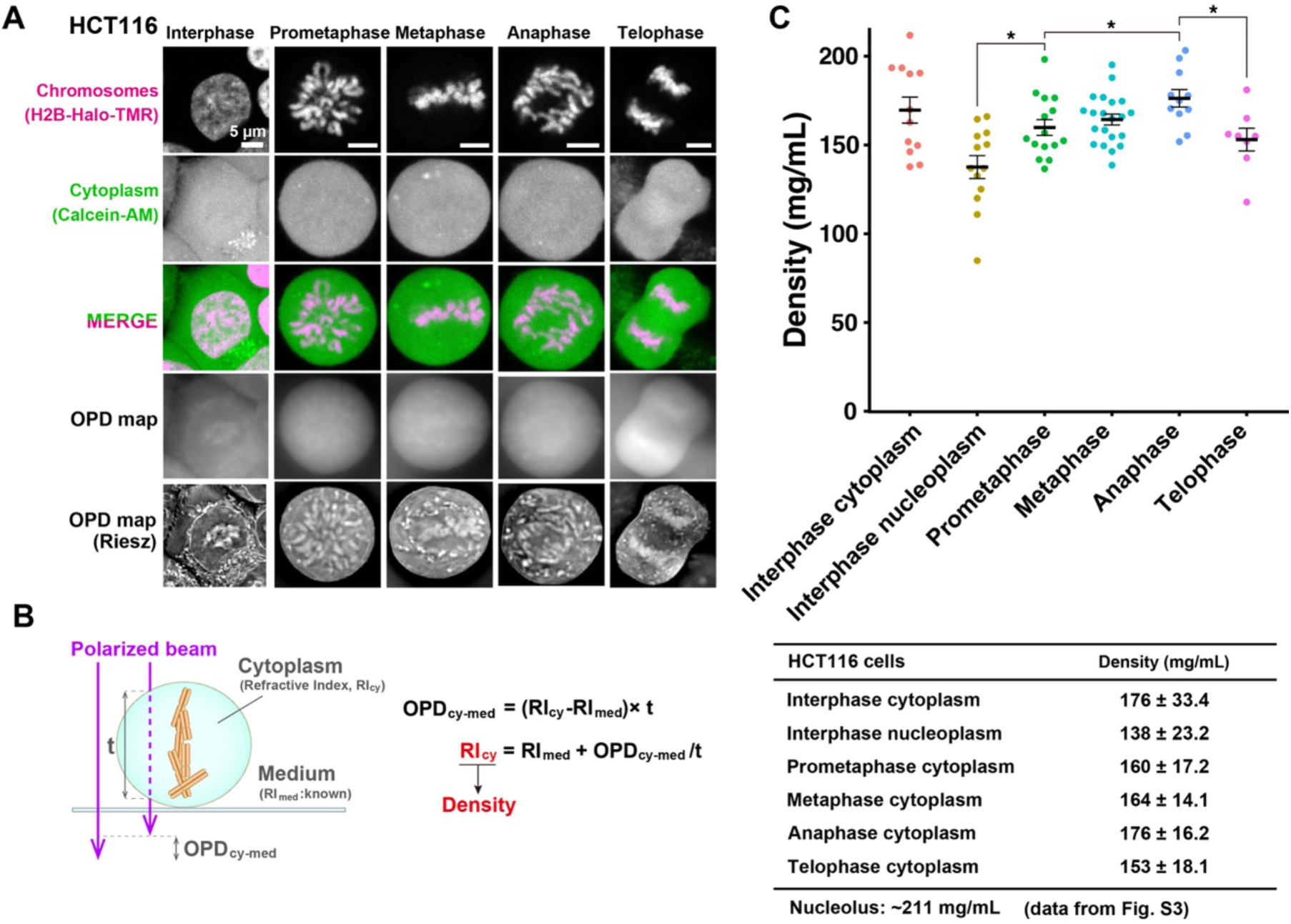
Density imaging of live human HCT116 cells by the new OI-DIC microscopy. (**A**) Confocal images of chromosome staining and cytoplasm staining, and OPD map and Riesz images obtained from OI-DIC imaging in live HCT116 cells. Riesz images are edge-enhanced OPD maps according to the inverse Riesz transform. To stain chromosomes, histone H2B-HaloTag was expressed in HCT116 cells and labeled with HaloTag ligand TMR. Cytoplasm was stained by Calcein-AM. Note that image intensities are not presented on the same scale for better visualization. Scale bar: 5 μm. **(B)** A simple schematic of the method to estimate the RI of cytoplasm (RI_cy_). The RI_cy_ can be calculated using the formula shown based on the OPD measured by OI-DIC, the RI of medium (RI_med_) measured by the refractometer, and thickness (t) values. For details, see SI Materials and Methods. **(C)** Total molecular densities of chromosome milieu gradually increased with the progression from prometaphase to anaphase in live HCT116 cells. Each dot represents the average of the estimated total density at several points within a single cell, where no apparent structure is apparent in the Riesz image (e.g., Fig. S3). For details, see SI Materials and Methods. Black bars show the mean, error bars indicate the SE, and cell number are N = 13 (Interphase cytoplasm), N = 13 (Interphase nucleoplasm), N = 15 (Prometaphase cytoplasm), N = 21 (Metaphase cytoplasm), N = 11 (Anaphase cytoplasm), and N = 8 (Telophase cytoplasm). *, P < 0.05 by the two-sided unpaired t-test for Interphase nucleus vs. Prometaphase cytoplasm (P = 9.5 × 10^−3^), for Prometaphase cytoplasm vs. Anaphase cytoplasm (P = 1.9 × 10^−2^), for Anaphase cytoplasm vs. Telophase cytoplasm (P = 1.2 × 10^−2^).

Based on the OPDs, thicknesses (t) of cytoplasm and nucleoplasm, and the known RI of the medium (1.3362 for McCoy’s 5A including 10% FBS in Fig. 1D(ii)), we calculated the RIs of cytoplasm (Figs. 2B and S2A) and nucleoplasm (Fig. S2B) in HCT116 cells (Table S1). It is known that there is a linear correlation between the concentrations of various macromolecules (proteins, lipids, carbohydrates, and nucleic acids) and the RIs of their solutions, within the physiological concentration range (40, 45, 46). According to the calibration curve of RIs versus the density of standard solutions for protein or nucleic acid (Fig. 1D(iii)), we obtained total density values in the cytoplasm and nucleoplasm of interphase cells and cytoplasmic density values of various stages of mitotic cells after nuclear envelope breakdown (NEBD) (Fig. 2C). Mitotic stages were judged by their chromosome morphology (Fig. 2A). Also note that we did not look at mitotic chromosomes themselves this time (Fig. 2B) since we wanted to examine molecular crowding around chromosomes, that is, the chromosome milieu.

### Molecular crowding in mitotic chromosome milieu is higher than in interphase chromatin

Our quantification (Fig. 2C; Table S1) shows that the density of interphase nucleoplasm is 138 mg/mL, which agrees well with our previous report (40) and is lower than that of cytoplasm (176 mg/mL). Interestingly, the density of cytoplasm in prometaphase cells, which corresponds to chromosome milieu, is 160 mg/mL (Fig. 2C) and higher than that of interphase nucleoplasm. With mitotic progression from prometaphase to anaphase, density gradually increased from 160 mg/mL to 176 mg/mL (Fig. 2C). Maximal cytoplasmic density in anaphase is consistent with previous reports on maximal chromosome compaction in anaphase (47, 48). Once the cytokinesis and chromosome decondensation started at telophase (Fig. 2A), the density was reduced (Fig. 2C). These results suggest that depletion force in mitotic chromosome milieu is higher than in interphase nucleoplasm and increases with mitotic progression from prometaphase to anaphase.

To further support our findings in HCT116 cells that depletion force around mitotic chromosomes is higher than in interphase chromatin, we examined another cell line, Indian Muntjac DM cells (Fig. 3A). DM cells were derived from deer fibroblast cells and have very large mitotic chromosomes (49–51). Chromatin and cytoplasm were labeled with Hoechst 33342 and Calcein AM simultaneously (Fig. 3A) to obtain thickness information (t) of interphase cytoplasm (Fig. S2A) and nucleoplasm (Fig. S2B), and mitotic cytoplasm (Fig. 2B).

**Figure 3.**
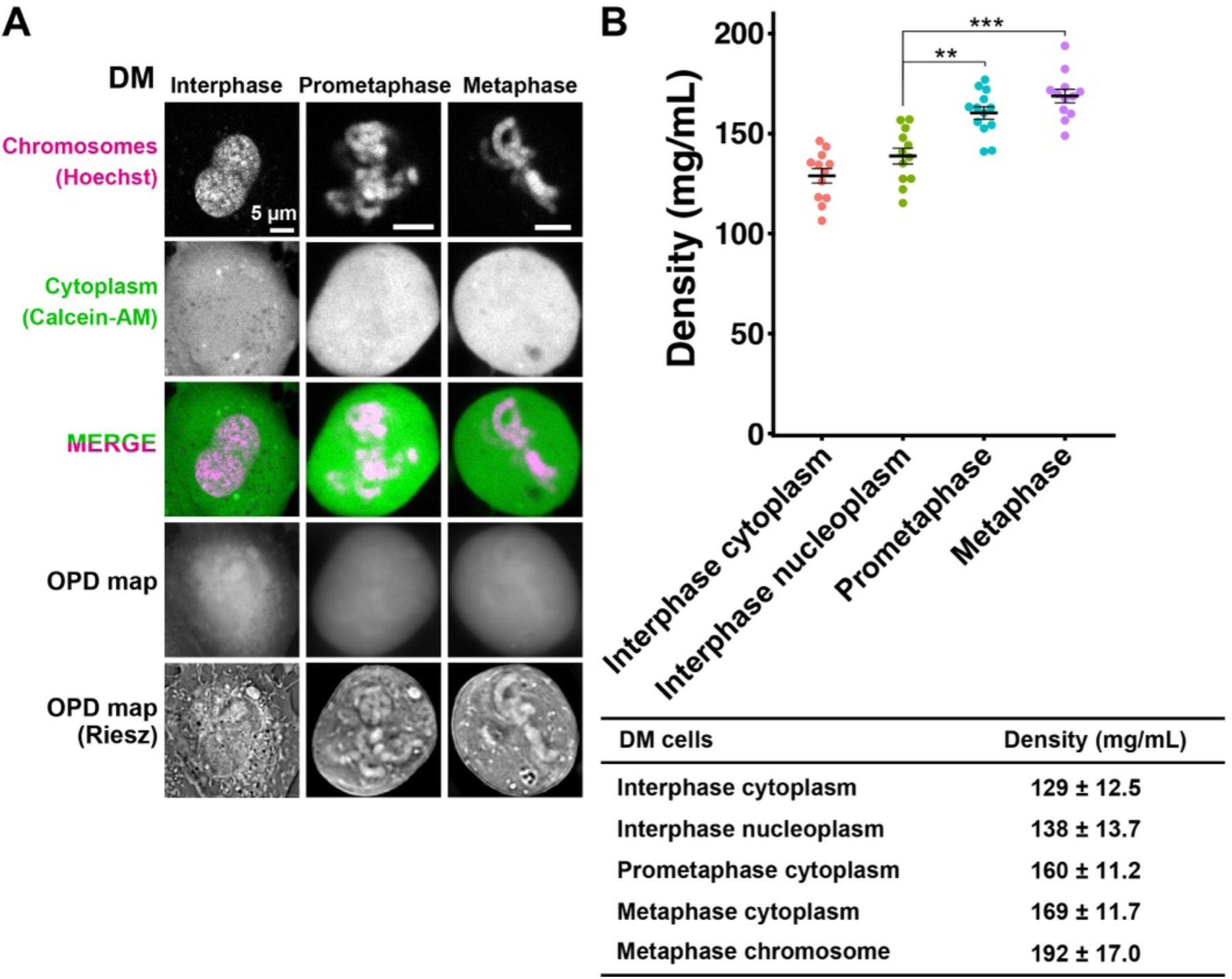
Density imaging of live Indian Muntjac DM cells by the new OI-DIC microscopy. (**A**) Confocal images of chromosome staining and cytoplasm staining, and OPD map and Riesz images obtained from OI-DIC imaging in live DM cells. Riesz images are edge-enhanced OPD maps according to the inverse Riesz transform. Chromosomes were stained with Hoechst 33342. Cytoplasm was stained by Calcein-AM. Note that image intensities are not presented on the same scale for better visualization. Scale bar: 5 μm. **(B)** Total molecular densities of chromosome milieu increased during mitosis in live DM cells. Each dot represents the average of the estimated total density at several points within a single cell, where no clear structure is apparent in the Riesz image. Black bars show the mean, error bars indicate the SE and cell numbers are N = 12 (Interphase cytoplasm), N = 12 (Interphase nucleoplasm), N = 13 (Prometaphase cytoplasm), and N = 12 (Metaphase cytoplasm). **, P < 0.001 by the two-sided unpaired t-test for Interphase nucleus vs Prometaphase cytoplasm (P = 3.2 × 10^−4^) and ***, P < 0.0001 for Interphase nucleus vs Metaphase cytoplasm (P = 9.4 × 10^−6^).

Using a similar procedure for HCT116 cells, we found that the density of cytoplasm in prometaphase DM cells, which corresponds to chromosome milieu, was 160 mg/mL (Fig. 3B) and higher than that of interphase nucleoplasm (138 mg/mL). Density increased from prometaphase to metaphase (Fig. 3B; Table S2). Obtained results in DM cells were consistent with those in human HCT116 cells, suggesting that a transient rise in the density of chromosome milieu may be a general feature of mitotic cells. These findings also suggest that depletion force can contribute to mitotic chromosome condensation.

### Density in mitotic chromosomes is 192 mg/mL

DM cells have much larger and fewer mitotic chromosomes (7-9 chromosomes/cell)(49–51) than HCT116 cells (∼46 chromosomes, relatively normal human karyotypes) (52). This difference enabled us to obtain the OPL of mitotic chromosomes (Fig. S5B) and measure the density in mitotic chromosomes as 192 mg/mL in DM cells (left, Fig. S6; Table S2). This value was significantly higher than that of mitotic cytoplasm (169 mg/mL, *p* = 0.0040) and comparable to the result of our previous study on the density (208 mg/mL) of the mouse pericentric heterochromatin (chromocenter) (40). However, the density of 192 mg/mL in DM mitotic chromosomes might be underestimated because it was difficult to accurately measure OPL within mitotic chromosomes, even when the chromosomes were larger (Fig. S5B and S5C). Also, note that the reported heterochromatin result was obtained using the earlier OI-DIC system that was not installed with CLSM (40), and the thickness (t) estimation of the target structures was not as accurate as the new OI-DIC system.

### Hypertonic treatment raised depletion force and induced chromosome hypercondensation

We wondered what would happen to the mitotic HCT116 cells and their chromosomes if we changed their cytoplasmic density. To address this question, we first performed the hypertonic treatment with 520 mOsm by adding concentrated phosphate-buffered saline (PBS) (Fig. 4A) as reported in (53, 54), while the physiological osmotic condition is 290 mOsm. Upon hypertonic treatment, the cytoplasmic density increased from 164 mg/mL to 228 mg/mL (Fig. 4B; Table S1). Mitotic chromosomes looked clumped together (Fig. 4A). The signal intensity of mitotic chromatin labeled by H2B-HaloTag-TMR also increased (Fig. 4C), suggesting a rise in depletion force and induced hypercondensation of mitotic chromosomes upon hypertonic treatment.

**Figure 4.**
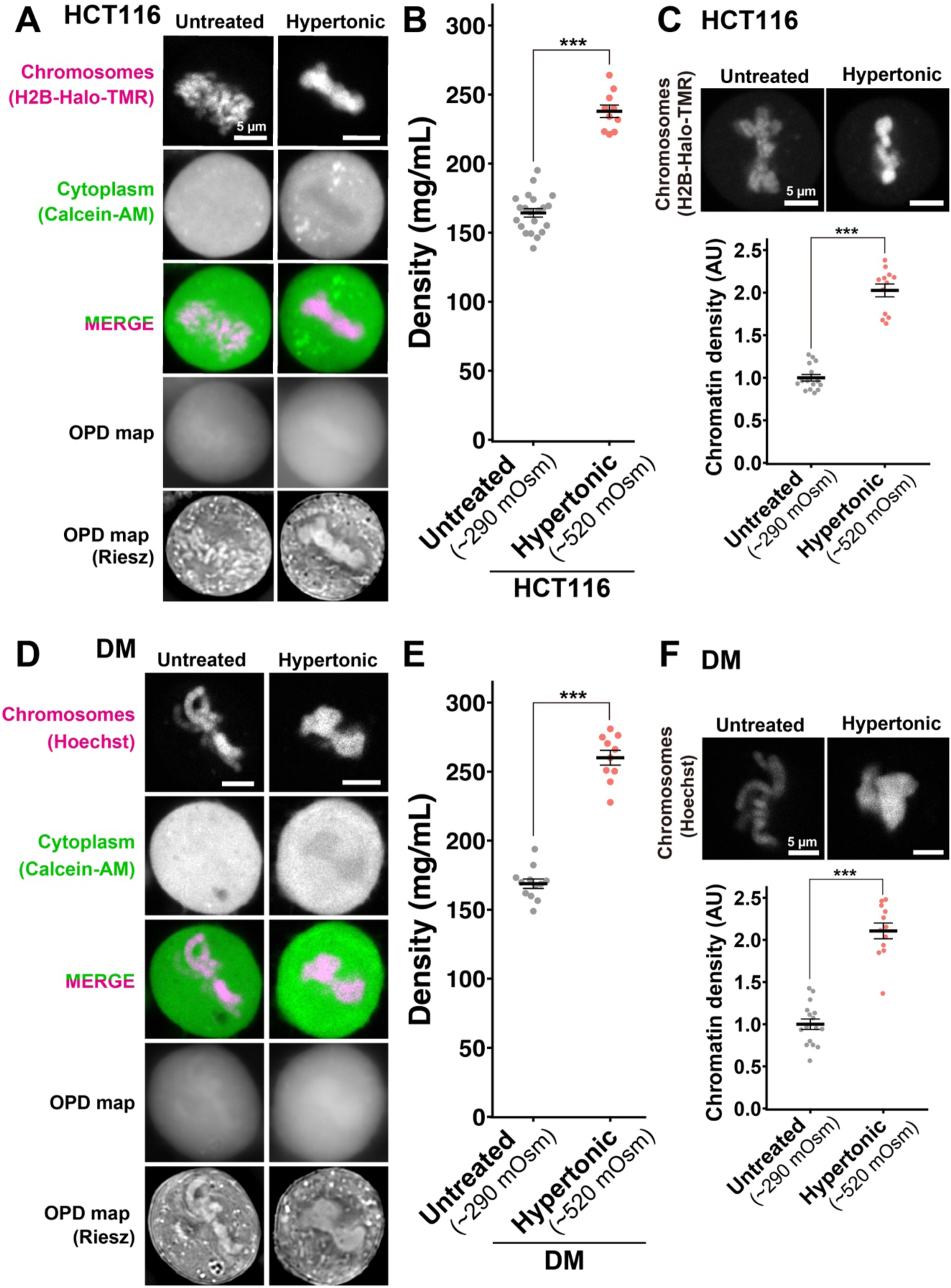
Density imaging of hypertonic mitotic cells by the new OI-DIC microscopy. (**A**) Confocal images of live HCT116 cells with hypertonic treatment, OPD map, and Riesz images obtained from OI-DIC imaging. Hypertonic treatment induced hypercondensation of chromosomes, making chromosomes fuse together. Chromosomes were labeled with histone H2B-HaloTag-TMR. Cytoplasm was stained by Calcein-AM. Note that image intensities are not presented on the same scale for visualization. Scale bar: 5 μm. **(B)** Hypertonic treatment increased the total densities of cytoplasm in mitotic live HCT116 cells. Each dot represents the average of the estimated total density at several points within a single cell, where no apparent structure is apparent in the Riesz image. Black bars show the mean, error bars represent the SE, and cell numbers are N = 21 (Untreated) and N = 10 (Hypertonic). ***, P < 0.0001 by the two-sided unpaired t-test (P = 1.0 × 10^−10^). Untreated data was reproduced from Fig. 2C (metaphase). **(C)** (Top) Hypertonic treatment induced hypercondensation of chromosomes of HCT116. Projection of 3 z-sections. Scale bar: 5 μm. (Bottom) Quantification of chromatin density in HCT116 cells with hypertonic treatment. Each dot represents the mean intensity of chromosome staining within a single cell, normalized by the mean intensity of untreated cells. Black dots show the mean, error bars represent the SE, and cell numbers are N = 15 (Untreated) and N = 15 (Hypertonic). ***, P < 0.0001 by Wilcoxon rank sum test (P = 1.3 × 10^−8^). AU, arbitrary units**. (D)** Confocal images of live DM cells with hypertonic treatment, OPD map, and Riesz images obtained from OI-DIC imaging. Hypertonic treatment induced hypercondensation of chromosomes, making chromosomes clump together. Chromosomes were stained with Hoechst 33342. Cytoplasm was stained by Calcein-AM. Note that image intensities are not presented on the same scale for visualization. Scale bar: 5 μm. **(E)** Hypertonic treatment increased the total densities of cytoplasm in mitotic live DM cells. Each dot represents the average of the estimated total density at several points within a single cell, where no clear structure is apparent in the Riesz image. Black dots show the mean, error bars represent the SE, and cell numbers are N = 12 (Untreated) and N = 10 (Hypertonic). ***, P < 0.0001 by the two-sided unpaired t-test (P = 2.1 × 10^−10^). Untreated data was reproduced from Fig. 2C (metaphase). **(F)** (Top) Hypertonic treatment induced hypercondensation of chromosomes in DM cells. Projection of 3 z-sections. Scale bar: 5 μm. (Bottom) Quantification of chromatin density in DM cells with hypertonic treatment. Each dot represents the mean intensity of DNA staining within a single cell, normalized by the mean intensity of untreated cells. Black bars show the mean, error bars represent the SE, and cell numbers are N = 16 (Untreated) and N = 16 (Hypertonic). ***, P < 0.0001 by Wilcoxon rank sum test (P = 1.3 × 10^−8^). AU, arbitrary units.

Similar findings were achieved using DM cells with hypertonic treatment (Fig. 4D). An increase in cytoplasmic density (Fig. 4E; Table S2) induced hypercondensation of mitotic chromosomes and chromosome clumping (Fig. 4D and 4F). In addition, the density within mitotic chromosomes increased from 192 mg/mL to 256 mg/mL (Fig. S6). The cells with a higher density in the cytoplasm tended to have a higher density for their chromosomes. These results support the hypothesis that depletion force contributes to mitotic chromosome condensation.

### Hypotonic treatment lowered depletion force and induced chromosome decondensation

Next, we treated HCT116 cells hypotonically (190 mOsm) by diluting the medium (Fig. 5A). We found that just after the treatment (<10 min), cytoplasmic density in prometa-/metaphase cells reduced from 164 mg/mL to 93 mg/mL (Fig. 5B; Table S1). Chromatin intensity labeled by H2B-HaloTag-TMR also decreased simultaneously (Fig. 5C), suggesting a considerable decondensation of mitotic chromosomes with a reduction in depletion force. These findings strengthen our notion that depletion force is involved in mitotic chromosome condensation.

**Figure 5.**
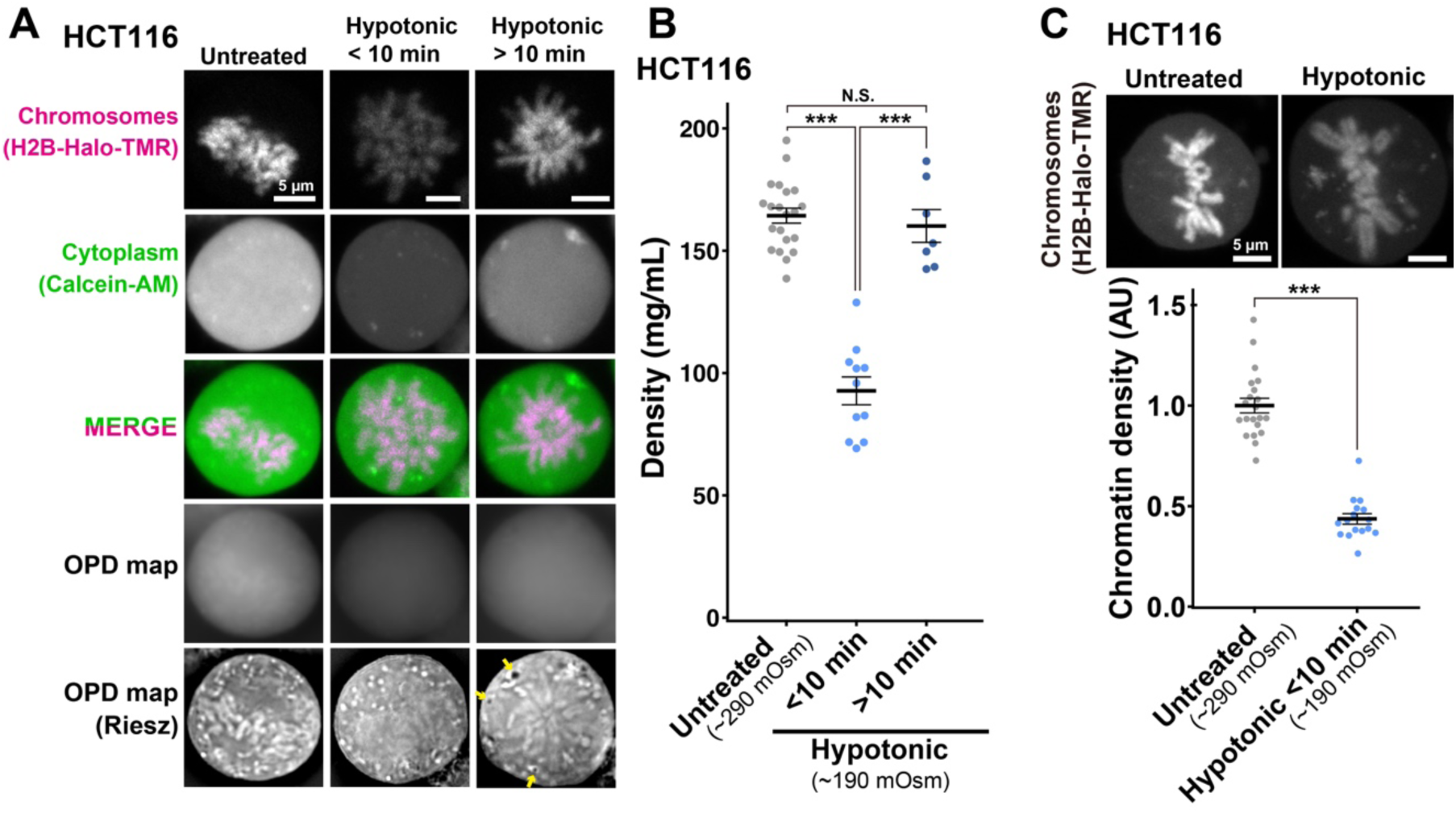
Density imaging of hypotonic mitotic HCT116 cells by OI-DIC microscopy. (**A**) Confocal images of live HCT116 cells with hypotonic treatment and OPD map and Riesz images obtained from OI-DIC imaging. Hypotonic treatment induced a transient decondensation of chromosomes (within 10 min after treatment), and then the chromosomes seemed to re-condense. Chromosomes were labeled with H2B-HaloTag-TMR. Cytoplasm was stained by Calcein-AM. Note that image intensities are not presented on the same scale for visualization. Scale bar: 5 μm. **(B)** Hypotonic treatment decreased the total densities of cytoplasm in mitotic live HCT116 cells. Each dot represents the average of the estimated total density at several points within a single cell, where no apparent structure is apparent in the Riesz image. Black bars show the mean, error bars represent the SE, and measured points are N = 21 (Untreated), N = 11 (Hypotonic < 10 min), and N = 7 (Hypotonic > 10 min). ***, P < 0.0001 by the two-sided unpaired t-test for Untreated vs Hypotonic < 10 min (P = 6.4 × 10^−9^) and for Hypotonic < 10 min vs Hypotonic > 10 min (P = 2.8 × 10^−6^). N.S., not significant (P = 0.58). Untreated data was reproduced from Fig. 2C (metaphase). **(C)** (Top) Hypotonic treatment (< 10 min) induced decondensation of chromosomes of HCT116. Projection of 3 z-sections. Scale bar: 5 μm. (Bottom) Quantification of chromatin density in HCT116 cells with hypotonic treatment (< 10 min). Each dot represents the mean intensity of DNA staining within a single cell, normalized by the mean intensity of untreated cells. Black dots show the mean, error bars represent the SE, and the cell numbers are N = 21 (Untreated) and N = 16 (Hypotonic). ***, P < 0.0001 by Wilcoxon rank sum test (P = 1.6 × 10^−10^). AU, arbitrary units.

We noticed that the rise in cytoplasmic density recovered >10 min after the hypotonic treatment (Fig. 5B), while cellular effects caused by hypertonic treatment seemed stable after 30 min, consistent with a previous report (55). The induced decondensation of chromosomes also attenuated on the same time scale (Fig. 5A). Interestingly, we found some low-density regions or vesicles in the cytoplasm of prometaphase cells appeared with prolonged hypotonic treatment (>10 min) (shown by arrows in Fig. 5A), suggesting the influx of water molecules were somehow stored and exported to keep intracellular osmotic pressure.

### Macromolecular crowders induce liquid droplets of chromatin *in vitro*, and additional crowding make the droplets stiffer and more solid-like

To finally examine how an increase in depletion force/macromolecular crowding during mitosis can change the physical properties of chromosomes, we performed an *in vitro* condensation assay using chicken native chromatin. The peak size of the native chromatin used for this assay was ∼6 kb and corresponded to ∼30 nucleosomes (Fig. S7A, B). Cations diminish the repulsion between negatively charged nucleosomes in the chromatin and electrostatically attract them to form chromatin condensates (19–23, 56) (25, 54, 57). Therefore, we incubated our native chromatin in a buffer containing 100 mM K^+^ and 0.8 mM Mg^2+^ to mimic physiological concentrations of cations in the cell (25) and found many fiber-like condensates of chromatin were formed (Fig. 6A, Panel 1), consistent with previous reports (25, 54, 57).

**Figure 6.**
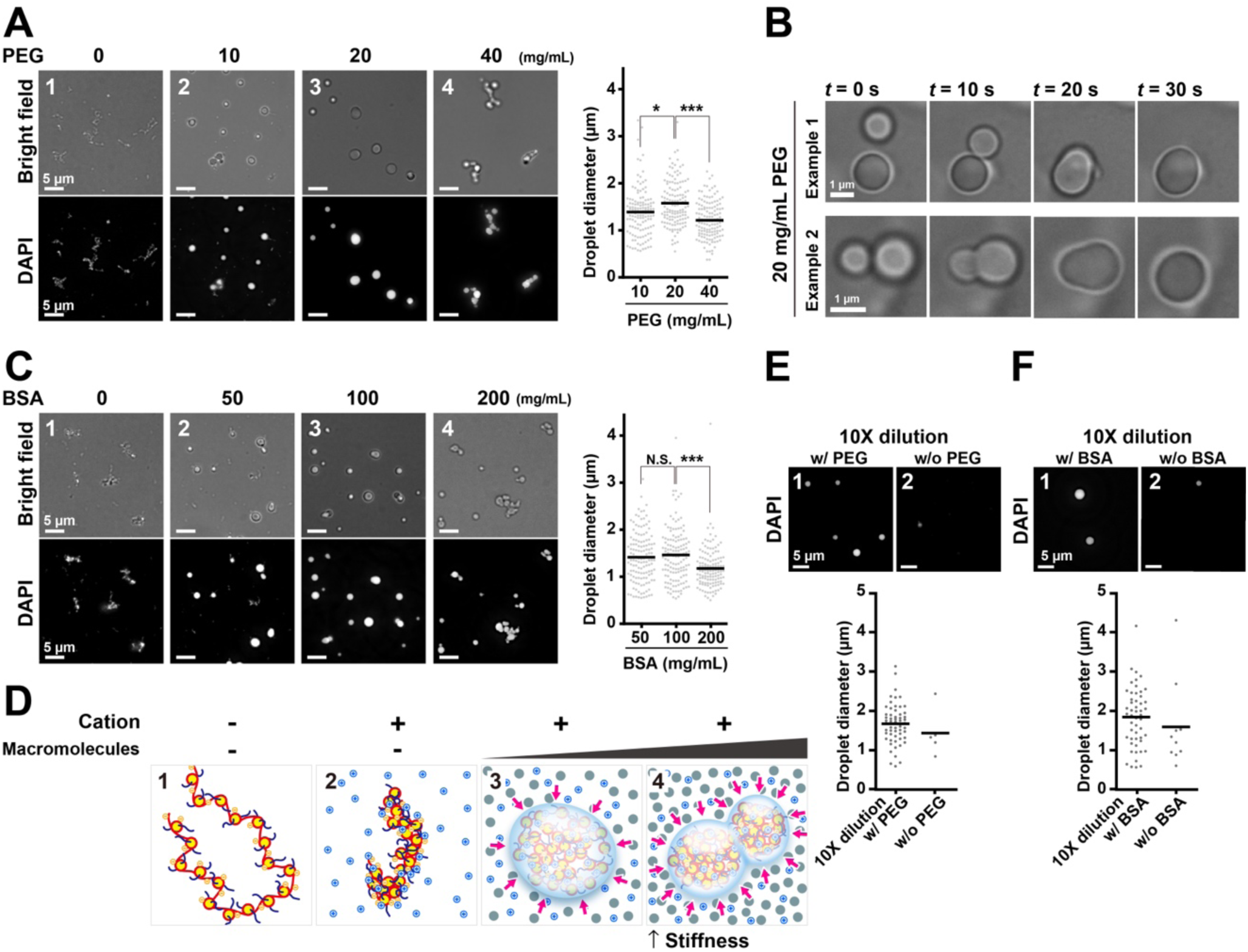
Liquid droplet formation and solidification of chicken native chromatin by macromolecular crowders. (**A**) Fibrous chromatin condensates were induced by 100 mM K^+^ and 0.8 mM Mg^2+^ (Panel 1, 0 mg/mL PEG). PEG converted the fibrous condensates into droplets (Panels 2 and 3). Further increases in the concentration of PEG put the droplets together without fusions, suggesting the solid-like property of the droplets (Panel 4). Diameters of formed droplets were shown on the right plots. *, P < 0.05 by Wilcoxon rank sum test for 10 mg/mL vs. 20 mg/mL PEG (P = 0.011). ***, P < 0.0001 for 20 mg/mL vs. 40 mg/mL PEG (P = 7.5 × 10^−9^). **(B)** Fusions of formed droplets. Two typical examples are shown. Also see Movie S1. **(C)** Similar analyses to (A) on BSA-induced liquid droplets. N.S., not significant by Wilcoxon rank sum test for 50 mg/mL vs. 100 mg/mL BSA (P = 0.78). ***, P < 0.0001 for 100 mg/mL vs. 200 mg/mL BSA (P = 7.0 × 10^−6^). **(D)** Schematics of conversion from fibrous chromatin condensates to liquid droplets. The chromatin is stretched by electrostatic repulsion without cations (Panel 1). Mg^2+^ and K^+^ decrease net negative charge and repulsion and induce fibrous chromatin condensates (Panel 2). Macromolecular crowding/ depletion force converts the fibrous condensates into droplets (Panel 3). Further crowding makes the droplets stiffer (Panel 4). **(E)** The droplet dilution assay. After the droplets were formed with 20 mg/mL of PEG, the reaction mixtures were diluted 10-fold in a buffer with (Panel 1) or without (Panel 2) 20 mg/mL of PEG. The images depict fewer and smaller droplets in the diluted buffer without PEG, suggesting that the droplets were dissolved in the diluted buffer. (Bottom) A quantitative analysis of the droplet dilution assay. The number and diameters of the droplets in a randomly picked area (6.0 x 10^4^ µm^2^) are plotted. **(F)** The dilution assay on BSA-formed droplets.

We simulated depletion force/macromolecular crowding of chromosome milieu by incubating increasing concentrations of polyethylene glycol (a neutral polymer; PEG, M.W. ∼8 kDa), bovine serum albumin (BSA, M.W. 66 kDa), or Dextran (M.W. ∼200kDa) with our fiber-like condensates of chromatin. We first added increasing concentrations of PEG as a crowder and observed morphological changes of chromatin condensates. With 10 mg/mL (w/v) (corresponding to 1.25 mM) PEG, some spherical structures were found among fibrous condensates (Fig. 6A, Panel 2). At 20 mg/mL (2.5 mM) PEG, almost all condensates became spherical with various sizes and looked like liquid droplets formed by liquid-liquid phase separation (LLPS) (Fig. 6A, Panel 3) (56, 58–60). Indeed, the droplets often fused to form larger droplets (Fig. 6B; Movie S1). Upon increase in PEG to 40 mg/mL (5.0 mM), droplets were connected to one another (Fig. 6A, Panel 4). Note that their sizes were not as large as those with 20 mg/mL PEG (Fig. 6A, right). These were presumably stuck together and not fused into lager droplets during our observation period (up to 1 h). Results using a higher concentration of PEG suggested further condensation of chromatin creates stiffer and more solid-like droplets, reducing their fluidity (Fig. 6D).

Next, we used BSA as another crowder in the same concentration of cations and native chromatin. BSA affected chromatin morphology at higher concentrations (w/v) than PEG. At 50 mg/mL (0.75 mM) BSA, chromatin droplets formed among fibers (Fig. 6C, Panel 2). Almost all condensates became droplets at 100 mg/mL (1.5 mM) BSA (Fig. 6C, Panel 3). As in the case of PEG, increasing the BSA concentration to 200 mg/mL (3.0 mM) formed clusters of droplets that appeared to stick together (Fig. 6C, Panel 4). Again, their sizes were smaller than those with 100 mg/mL (1.5 mM) BSA due to much fewer fusion events being prevented by stiffer and more solid-like properties (Fig. 6C, right). We also obtained a similar set of results using Dextran (M.W. ∼200kDa) (Fig. S7C). These results suggest that depletion force/macromolecular crowding can change the physical properties of chromosomes. Addition of crowders induced the formation of chromatin liquid droplets *in vitro*. Further crowding pushed chromatin droplets from the outside, condensed chromatin, and increased droplet stiffness, which may have prevented droplet fusions at our time scale (up to 1 h) due to the reduced fluidity (Fig. 6D).

To examine whether the chromatin droplet formation induced by depletion force/macromolecular crowding was reversible, we lowered the crowding effect around the droplets. After droplets were made using 20 mg/mL PEG, 100 mg/mL BSA, or 50 mg/mL Dextran, the droplet solutions were diluted 10-fold with a buffer containing 100 mM K^+^ and 0.8 mM Mg^2+^ with or without crowder (Figs. 6E, F and S7D). The number of droplets in the diluted buffer without crowder substantially decreased and more small chromatin condensates appeared as compared to the control droplets in the diluted buffer with crowder. Droplets resolved into small chromatin condensates, suggesting that droplet formation by depletion force/macromolecular crowding is reversible.

## Discussion

The chromosome milieu governs physical forces contributing to chromosome condensation. We developed a novel OI-DIC microscopy system combined with CLSM to elucidate the effects of the chromosome milieu in live mitotic cells (Fig. 1B). The subsequent analysis enabled the absolute density in cytoplasm of various stages of mitotic human cells to be calculated, as well as in the cytoplasm and nucleoplasm of interphase cells. We demonstrated that the density of cytoplasm in metaphase cells (164 mg/mL) was higher than that of interphase nucleoplasm (138 mg/mL). Density gradually increased from 160 mg/mL to 176 mg/mL (Fig. 2C) as cells mitotically progressed from prometaphase to anaphase, concurring with chromosome condensation (47, 48). Furthermore, hypertonic treatment of the mitotic cells rapidly raised density and induced chromosome condensation, while hypotonic treatment had the opposite effects. Our findings suggest that a transient rise in depletion force (the density of proteins, RNAs, and others) during mitosis is involved in mitotic chromosome condensation, providing a novel insight into the physical bases of mitotic chromosome condensation in living human cells.

To our knowledge, we are the first to demonstrate that the depletion force/macromolecular crowding converts fibrous chromatin condensates formed by cations into liquid droplets *in vitro*. Previous reports on chromatin liquid droplet formation have thus far only used synthetic nucleosomes such as 12-mer arrays, which have uniform properties in the length, spacing, size, and modifications (54, 56, 61). The liquid droplets we observed using native chromatin reminded us of the chromatin liquid droplets in living mitotic cells when Schneider et al. injected cells with the restriction enzyme AluI to fragment chromatin (15). Increasing depletion force/macromolecular crowding *in vitro* made the droplets stiffer and more solid-like, with further chromatin condensation (Fig. 6D). While cations induced an electrostatic attraction force between chromatin, the depletion force/macromolecular crowding worked as external pressure from outside (Fig. 6D). Our results suggest that the depletion force/macromolecular crowding is another force that contributes to chromosome rigidity during mitosis (Fig. 7). The additional condensation by increased depletion force is particularly advantageous for the chromosome segregation and transmission processes during anaphase, when mechanical shearing stress is high. Note that the highest cytoplasmic density level was identified in anaphase (Fig. 2C) and concurred with previous reports of the highest level of chromosome compaction in anaphase (47, 48). Condensed chromatin domains in interphase have been observed in a variety of cells (62–66) and are proposed to work as building blocks for mitotic chromosomes (5, 64). The assembled stiffer chromatin droplets that we observed in the highly crowding situation looked like mitotic chromosomes (Panels 4 in Fig. 6A and C and Fig. S7C). Hence, the depletion force/macromolecular crowding may facilitate the assembly of condensed chromatin domains to form mitotic chromosomes.

**Figure 7.**
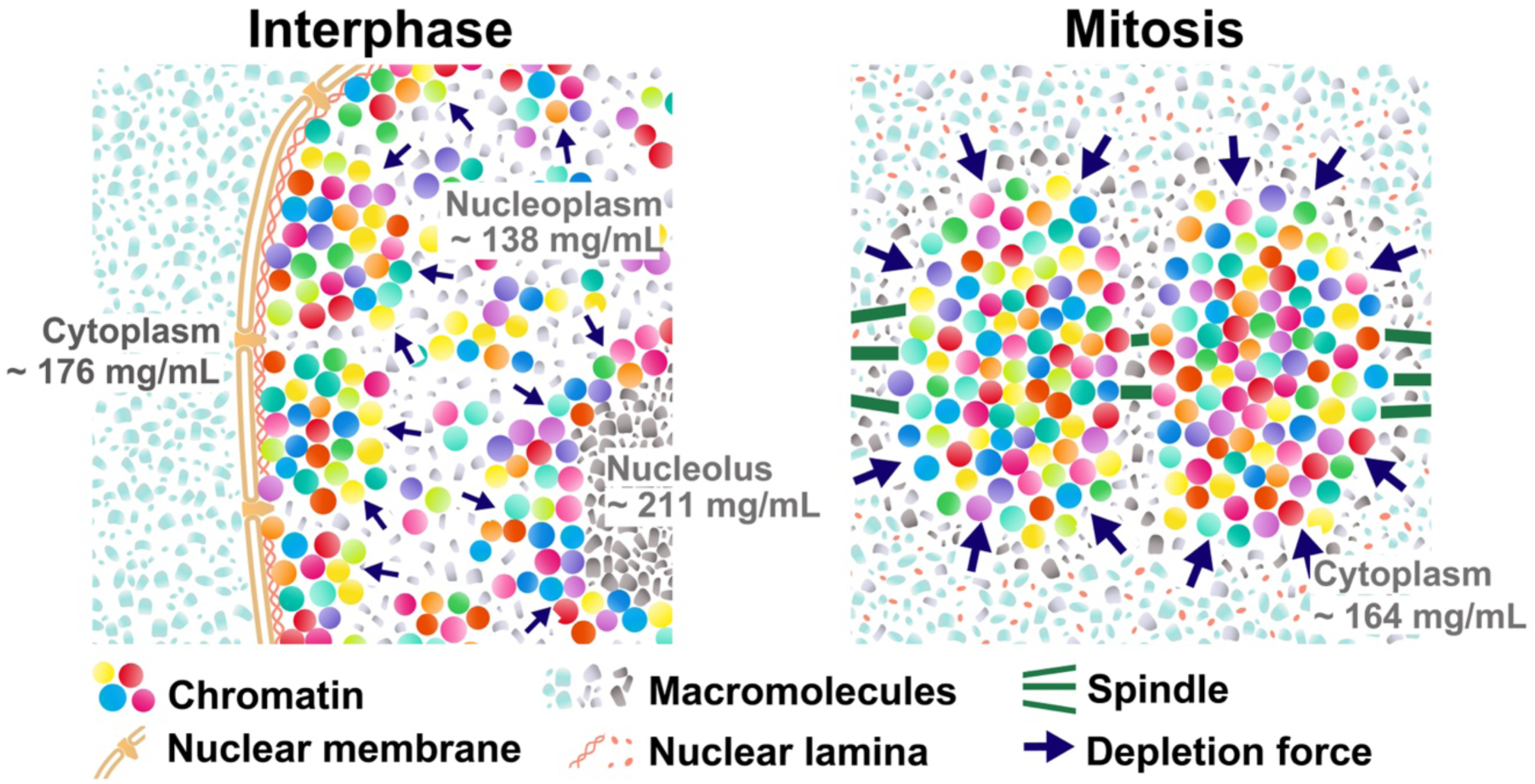
Model for upregulation of macromolecular crowding in mitotic cells. (Left) Schematic of macromolecular crowding in interphase live cells with soluble macromolecules in cytoplasm (light-blue), nucleoplasm (gray), and nucleolus (dark gray). The cytoplasm, nucleus, and nucleolus are compartmentalized and not mixed during interphase. Molecular densities of nucleoplasm are lower than the cytoplasm and nucleolus. (Right) After NEBD, soluble macromolecules that were localized to the cytoplasm, nucleus, and nucleolus at interphase are now mixed. Molecular density of chromosome milieu increases, making the macromolecular crowding effect stronger, and contributes to local condensation of chromosomes. Note that these schematics are highly simplified models and also that depletion force work even in interphase chromatin (small navy arrows).

What is the underlying mechanism of upregulating macromolecule density during mitosis? Small molecules (<40-60 kDa) can path through the nuclear pores by passive diffusion (67) and are rather evenly distributed within the cell. On the other hand, large macromolecules and their complexes, including RNAs, DNAs, and lipids, in eukaryotic cells are highly compartmentalized in interphase by nuclear membrane (left, Fig. 7; Fig. 2A). We propose that cells regulate their density localizations by compartmentalization and can upregulate cytoplasmic density after NEBD (right, Fig. 7). OPD maps of interphase cells show that cytoplasmic organelles (ER, Golgi apparatus, mitochondria, etc.), the nuclear membrane structure, and nucleoli are prominent high-density structures (bottom left, Fig. 2A). Note that we avoided such cytoplasmic organelles for our measurements (Fig. S3).

Upon NEBD, the nuclear envelope, nuclear pore complexes, nuclear lamina, and nucleoli are disassembled into small pieces (complexes or vesicles) for cell division and become a part of the “chromosome milieu.” As a result, cytoplasmic and nucleolus factors are exposed to chromosomes and fully contribute to an increase in depletion force/macromolecular crowding (right, Fig. 7). For instance, nucleoli, which are enriched with proteins and RNAs, occupy ∼12% of the nuclear volume and have a high density of >200 mg/mL (Fig. S4; (40)). Ribosomes are also a potential contributor to increase depletion force/macromolecular crowding during mitosis. We indeed observed that in interphase and prophase, prior to NEBD, ribosome components were sequestered from chromatin and confined to the cytoplasm by the nuclear membrane. In contrast, in prometaphase and metaphase, after NEBD, they can come into contact with mitotic chromosomes (Fig. S8). In addition, some of the nucleolar components also locate to the chromosome periphery, possibly causing additional depletion force (e.g., (68–70)). Together, these fully coordinated mitotic events can lead to a transient rise in depletion force/macromolecular crowding observed during mitosis (right, Fig. 7).

A transient rise in depletion force during mitosis can act as an additional force to form larger chromosomes in higher eukaryotic cells. Higher eukaryotic cells have a mitotic process called open mitosis. NEBD occurs during prophase, and the nuclear envelope is regenerated around the chromosomes in telophase (6, 7). Our findings suggest that the open mitosis system allows the cell to upregulate depletion force during mitosis. Closed mitosis occurs in the case of lower eukaryotes (e.g., fungi and yeasts), where the nuclear envelope does not break down during mitosis (71, 72). Their “chromosome milieu” seems constant during mitosis, suggesting that eukaryotic organisms might have evolved to acquire open mitosis and an additional chromosome compaction force for their larger chromosomes. As discussed above, this force could be beneficial to make larger chromosomes more rigid to ensure faithful transmission into daughter cells.

Our model described above (Fig. 7) is well compatible with reports by Son et al. (73) and Zlotek-Zlotkiewicz et al. (74), who measured the volume and density dynamics in proliferating mammalian cells. They found a rapid increase in cell volume and a decrease in whole cell density just as cells entered prophase. This prophase cell swelling seems to be driven by an osmotic water exchange (73), possibly to generate a larger, rounder space to promote accurate and rapid chromosome segregation in the context of animal tissue (73). Subsequently, cell volume decreased in the metaphase-anaphase transition. The reduction of cell volume can also contribute to an increase in molecular density with the mitotic progression from prometaphase to anaphase (Fig. 2C), promoting further chromosome condensation during mitosis, as shown in (47, 48, 75). Furthermore, our finding is consistent with data from an intracellular crowding sensor GimRET (76). A transient rise in the GimRET signal during cell division of HeLa cells (Fig. S8 in) suggested an up-regulation of depletion force during the chromosome condensation process.

The new OI-DIC module combined with CLSM can obtain matched high-resolution OPD and confocal images to get precise absolute densities in live cells, while another density microscopy may have difficulty because it requires a separate reference beam and uses two objective lenses for OPD/RI reconstruction. The new OI-DIC imaging system offers fresh insight into the density of live cellular environments, which govern the functions of protein/RNA factors and their complexes via their diffusions.

## Materials and Methods

For information on OI-DIC/confocal setup/imaging, cell lines, density estimation of cellular contents, chromatin preparation, and condensation/droplet formation assay, please refer to SI Appendix, Materials and Methods.

## Acknowledgments

We are grateful to Dr. K.M. Marshall for critical reading and editing of this manuscript and Dr. R. Imai for a preliminary result on density in mitotic chromosomes. We thank Dr. M. Kanemaki for providing their HCT116 cell line, Dr. H. Kimura and Dr. P. Cook for providing their DM cell line, Dr. Uchiumi for providing anti-ribosomal P protein antibody, Dr. Y. Murayama, Dr. A. Nakano, Dr. K. Hibino, Mr. M. A. Shimazoe, and Maeshima laboratory members for their helpful discussions and support. This work was supported by JSPS grants JP21H02453 (K.M.), JP22H05606 (S.Ide), JP21H02535 (S.Ide), JP20H05936 (K.M.), JP23K17398 (K.M., S.Ide, T.T.), Takeda Science Foundation, Inoué Endowment Fund (M.S.) and NIGMS/NIH grant R01GM101701 (M.S.). S.Iida was a SOKENDAI Special Researcher (JST SPRING JPMJSP2104) and supported by a SOKENDAI Genetics Course travel fellowship. S.Iida is currently a JSPS Fellow (JP23KJ0996).

