## Supplemental Information for "Orientation-Independent-DIC imaging reveals that a transient rise in depletion force contributes to mitotic chromosome condensation"

**This PDF file includes:**

Materials and Methods
Figures S1 to S8
Tables S1 to S2
Legends for Movie S1
SI References

**Other supporting materials for this manuscript include the following:**

Movie S1

### Materials and Methods

#### Cell lines and establishment of stable cell lines.

HCT116 cells (CCL-247; ATCC) were cultured at 37 °C and 5% $CO_2$ in McCoy's 5A medium (SH30200.01; HyClone) supplemented with 10% fetal bovine serum (FBS; FB-1061/ 500; Biosera). Indian Muntjac DM cells were a generous gift from Dr. H. Kimura and Dr. P. Cook (Tokyo Tech and Oxford University, respectively) (1). The cells were cultured at 37 °C and 5% $CO_2$ in Dulbecco's Modified Eagle's medium (DMEM; D5796- 500ML; Sigma-Aldrich) supplemented with 20% FBS.

The transposon system was used to stably express H2B-Halo in the HCT116 cell line. The constructed plasmid pPB-CAG-IB-H2B-HaloTag (2) was cotransfected with pCMV-hyPBase (provided by Sanger Institute with a materials transfer agreement) into HCT116 cells with the Effectene transfection reagent kit. Transfected cells were then selected with 10 µg/mL blasticidin S. HCT116 cells expressing SMC2-mAID-mClover and Tet-OsTIR1 were used as parental cells (3, 4).

pPB-CAG-mCherry-NPM1 plasmid was constructed on the basis of pPB-CAG-IB (provided from Sanger Institute with a materials transfer agreement). Full length NPM1 fused to EGFP at the N-terminus was cloned into pEF1-FRT plasmid (5). From pEF1-EGFP-NPM1-FRT, the coding region of NPM1 was amplified using the following primer pairs: 5'-GACGAGCTGTACAAGGGTGGAGGTGGATCTGGTGGAG-3' and 5'-GGGGCGGAATTCGTTTTAAAGAGACTTCCTCCACTGCCAG-3'. The mCherry fragment was amplified from pmCherry_a_tubulin_IRES_puro2 (#21043, Addgene) using the following primers: 5'- GAATTGATCTCTCGAGATGGTGAGCAAGGGCGAGGAG-3' and 5'-CTTGTACAGCTCGTCCATGCC-3'. The amplified fragments were joined together using standard overlapping PCR and inserted into pPB-CAG-IB-H2B-HaloTag digested with XhoI and HpaI using In-Fusion (639650; Takara). To stably express mCherry-NPM1 in HCT116 cell line, the transposon system was used. The constructed plasmid pPB-CAG-mCherry-NPM1 was cotransfected with pCMV-hyPBase into HCT116 cells expressing mAID-mClover-RPA194 (6) with the Neon Electroporation System (MPK5000; Thermo Fisher Scientific). For the electroporation, cells were resuspended in Neon Resuspension Buffer R (10 µL, Neon kit, Invitrogen) to a final concentration of $2.5 \times 10^7$ cells/mL and mixed with 500 ng of pPB-CAG-mCherry-NPM1 and pCMV-hyPBase, respectively. The mixture was pulsed once with a voltage of 1400 and a width of 20. Transfected cells were then selected with 10 µg/mL blasticidin S (029–18701; Wako).

#### Description of OI-DIC/confocal setup.

A simplified optical schematic of the confocal laser scanning microscope Olympus FV3000 (Evident, Tokyo, Japan) with the OI-DIC module is shown in Fig. 1B. The FV3000 consists of an inverted microscope Olympus IX83, a confocal scan unit, and a laser combiner. The microscope IX83 was equipped with a 100-W halogen lamp U-LH100L-3, a bandpass filter 546/10 nm (Chroma, Bellows Falls, VT, USA), a condenser IX2-DICD with front lens U-TLD 0.9NA, a water immersion objective lens UPlanSApo 60x/1.2NA W, and an 8-positions mirror turret FV30-RFACA. The condenser, objective lens, and specimen under investigation were placed in a Thermo Box IX83TB (Tokai Hit, Shizuoka, Japan). The turret contains an LSM mirror and analyzer cube IX3-FDICT. Regular Olympus DIC sliders were replaced by custom beam-shearing DIC assemblies. The first assembly consists of a pair of DIC prisms U-ODIC100HR, a liquid crystal polarization rotator, and a liquid crystal variable retarder (ARCoptix, Neuchatel, Switzerland). The second assembly consists of a pair of DIC prisms U-DICTHR and a liquid crystal polarization rotator. The assemblies can quickly switch the shear direction by 90º and change the bias without any mechanical movement. A set of 4 or 6 DIC images was captured by a digital camera Infinity 3-1M (Lumenera, Ottawa, Canada). More details about OI-DIC image acquisition and processing were described previously (7, 8). The laser combiner has four laser diodes with wavelengths of 405 nm, 488 nm, 562 nm, and 640 nm. Their beam intensities were controlled by an acousto-optic tunable filter and then delivered by a single-mode optical fiber to a confocal scan unit. Excitation dichroic mirror reflects the laser light to a pair of Galvano scanning mirrors and CLSM mirror. The emission light is transmitted by the dichroic mirror to an adjustable confocal pinhole and 2-channel multi-alkali PMT spectral detector unit FV31-SD. The spectral detector employs efficient Volume Phase Holographic transmission grating and adjustable slit with 1–100 nm bandwidth from a 400–800 nm detection region.

**Mathematical model of OI-DIC microscopy**

For the convenience of the reader, we briefly explain the mathematical principles of the OI-DIC technique and one of image processing algorithms. The OI-DIC mathematical model, along with several processing algorithms, were reported in detail elsewhere (8-11).

The intensity distribution in the DIC image $I(x,y)$ can be described by using a model of interference of two overlapping identical coherent images with an optical path difference $OPD(x,y)$, slightly offset from each other:

$$I(x,y) = \tilde{I}\sin^2\left\{\frac{\pi}{\lambda}\left[\Gamma + \mathbf{d}\cdot\mathbf{G}(x,y)\right]\right\} + I_c(x,y)$$

,

where $\tilde{I}$ is the initial beam intensity, $\lambda$ is wavelength, $\Gamma$ is bias, $\boldsymbol{d}$ is shear vector, **G**(x,y) is the optical path difference gradient vector, and $I_c(x,y)$ corresponds to an offset of the intensity signal, which is caused by the stray light.

The optical path difference gradient vector **G**(x,y) is the following:

$$\mathbf{G}(x,y)=\left(\frac{d(OPD(x,y))}{dx},\frac{d(OPD(x,y))}{dy}\right)=\left(\gamma(x,y)\cos\theta(x,y),\gamma(x,y)\sin\theta(x,y)\right),$$

where $\gamma(x,y)$ and $\theta(x,y)$ are the optical path difference gradient magnitude and azimuth, respectively.

In order to map the optical path difference gradient vector **G**(x,y), the OI-DIC microscope varies shear vector $\boldsymbol{d}$ and bias $\Gamma$. We capture two sets of raw DIC images at shear directions 0º (X-shear) and 90º (Y-shear) with negative, zero, and positive biases: $-\Gamma_0$, 0, and $+\Gamma_0$. Typically, we use biases of ±0.15λ and 0.

The following group of equations represents these six DIC images:

$$I_{i,j}(x,y)=\tilde{I}\cdot\sin^2\left\{\frac{\pi}{\lambda}[j\cdot\Gamma_0-d\cdot\gamma(x,y)\sin(\theta(x,y)-i\cdot 90°)]\right\}+I_{min}(x,y),$$

where $i$=1 and $i$=2 represent X- and Y-shears, respectively, $j$ = -1, 0, 1 corresponds to three settings of the bias, $d$ is the shear amount (magnitude of shear vector), and $I_{min}(x,y)$ corresponds to an offset of the intensity signal, which is caused by the stray light.

Initially two terms are computed ($i$ = 1, 2):

$$A_i(x,y)=\frac{I_{i,1}(x,y)-I_{i,-1}(x,y)}{I_{i,1}(x,y)+I_{i,-1}(x,y)-2I_{i,0}(x,y)}\tan\left(\frac{\pi\Gamma_0}{\lambda}\right).$$

Using the obtained terms, we can calculate the quantitative two-dimensional distributions of the gradient magnitude and azimuth of an optical path difference in the specimen as:

$$\gamma(x,y)=\frac{\lambda}{2\pi d}\sqrt{\sum_{i=1}^{2}arctan^2[A_i(x,y)]},$$

$$\theta(x,y)=arctan\left[\frac{\arctan(A_2(x,y))}{\arctan(A_1(x,y))}\right].$$

The gradient magnitude represents an increment of the optical path difference, which is in nanometers, along a lateral coordinate, which is also in nanometers. Thus, the gradient magnitude is unitless.

The obtained two-dimensional distribution of the optical path difference gradient vector **G**(x,y) is used for computing the optical path difference map *OPD(x,y)*. For this purpose, the gradient vector **G**(x,y) can be presented as a complex number:

$$\mathbf{G}(x,y) = \frac{\partial(OPD(x,y))}{\partial x} + i\frac{\partial\phi(OPD(x,y))}{\partial y} = \gamma(x,y)e^{i\theta(x,y)}, \quad (1)$$

where real and imaginary parts are X- and Y- components of the gradient vector, respectively.

At first, we apply the 2-dimensional Fourier transform to the left and middle parts of the equation above. The resultant integral equation can be solved by partial integration. After using the inverse 2-dimensional Fourier transform we receive the following formula for computation of the optical path difference *OPD(x,y)*:

$$OPD(x,y) = F^{-1}\left[\frac{F\left[\mathbf{G}(x,y)\right]}{i\left(\omega_x + i\omega_y\right)}\right],$$

where $\omega_x$ and $\omega_y$ are spatial angular frequencies.

Considering the right part of equation (1), we finally obtain the formula for computing the OPD:

$$OPD(x,y) = F^{-1}\left[\frac{F\left[\gamma(x,y)e^{i\theta(x,y)}\right]}{i\left(\omega_x + i\omega_y\right)}\right].$$

Then a computed two-dimensional distribution of the optical path difference is transformed into quantitative 8-bit grayscale image (map), where the image brightness is linearly proportional to value of the OPD and the maximum gray level of 255 corresponds to the chosen OPD ceiling.

**OI-DIC imaging of glass rods.**

A small number of glass rods 4 μm in diameter were suspended in two types of mineral oil with refractive indices of 1.54 and 1.58. Approximately 2 μL of the suspended solution was sandwiched between a glass slide and a coverslip and then sealed with nail polish. The glass rods in the mineral oil were analyzed by OI-DIC microscopy using the same procedure as the live cell imaging. A plot of the intensity profile across a glass rod was created by ImageJ (line width: 10 pixels) to validate density imaging by OI-DIC microscopy. The theoretical OPD curve was calculated based on the thickness and the RI of a glass rod, assuming that the cross-section of the glass rod is a circle with a diameter of 4 μm.

**Measurement of the Refractive Index (RI) of standard solutions.**

To obtain calibration curves of the RIs of proteins and nucleic acids (Fig. 1D (iii)), bovine serum albumin (BSA; Sigma; A9647-100G) and salmon sperm DNA (WAKO; 043-31381) were dissolved in water at concentrations of 0–200 and 0–30 mg/mL, respectively. The RIs of the prepared standard solutions were measured with a refractometer, Abbé-3L (Bausch & Lomb). The measured RIs and solution densities were plotted and fitted with linear functions to obtain calibration curves. RI of the medium was also measured by the same refractometer. The RI of McCoy's 5A/10% FBS was 1.3362 and the RI of DMEM/20% FBS was 1.3379.

**Cell preparation.**

Cells were seeded on 24 mm × 24 mm square glass coverslips coated with 2.5 μg/mL fibronectin (354008; Corning) diluted in PBS for 1 h at 37 °C and cultured for 1‑2 days. In HCT116 cells, cytoplasm and histone H2B-Halo were labeled with 3 μg/mL Calcein-AM and 5 nM TMR, respectively, for 30 min at 37 °C in 5% $CO_2$, then washed with 1× HBSS (H1387; Sigma-Aldrich) three times. In DM cells, DNA was stained with 1.8 μg/mL Hoechst33342 (H342; Dojindo) instead of TMR. Cells were then incubated in the following phenol red-free media. HCT116 cells were observed in McCoy's 5A (1-18F23-1; BioConcept) with 10% FBS, and DM cells were observed in DMEM (21063-029; Thermo Fisher Scientific) with 20% FBS. The cells were mounted on a glass slide with a silicone spacer having a thickness of 0.5 mm. We observed the mounted cells by OI-DIC and fluorescence imaging. Cells were not synchronized.

**Live-cell OI-DIC microscopy imaging.**

Before the experiment, the Thermo Box was set at 37 °C. The specimen was placed on the microscope stage, and the microscope was switched to the DIC mode. The analyzer was in the beam path, and the LSM mirror was out. The camera showed a live DIC image. The objective lens was focused on the middle plane of the cell under investigation. Using the custom software, which controlled the beam-shearing assemblies, we captured a set of DIC images with X- and Y-shear directions. We took 3 images in each direction, with zero and ±0.15λ biases. The total acquisition time of six images was about 1 s. The corresponding background image without cells or structures was also taken prior to each image acquisition to create the OPD map. The captured images were processed according to the previously reported algorithm (8), and the OPD map was generated (7) (Fig. S1). In the next step, the microscope was switched to the confocal mode. The analyzer was out of the beam path, and the LSM mirror was in. Then, we captured the z-stack and found the thickness of the cell, t.

**Density estimation of cellular contents.**

This OI-DIC version allowed the OI-DIC image and confocal images to be obtained in the same field of view. Therefore, it was possible to estimate both the OPD and thickness at a particular point and obtain an accurate molecular density at that point. The OPD was proportional to the thickness of a sample and the difference in RI between the sample and the surrounding solution, as shown in Fig. 1D(i). Therefore, we calculated the RI of samples based on the RI of the surrounding solution and sample thickness.

First, we cropped the OPD map to manually match the field of the confocal image captured by CLSM using intracellular structures (e.g., nuclei) as indicators. Second, to calculate the Optical Path Difference (OPD) between the medium and samples, we obtained the Optical Path Length (OPL) of medium, $OPL_{med}$, and OPL of sample, $OPL_s$. The pixel intensity of OPD map reflects OPL at that location. To obtain $OPL_{med}$, we set 5 ROIs [circles with a diameter of ~0.6 μm] outside of cells and calculated the mean intensity inside the ROIs. For $OPL_s$, we placed the ROIs at the several selected points within the sample. When calculating cytoplasmic OPL ($OPL_{cy}$), the ROIs were set to avoid apparent structures in the Riesz image and chromosomes in the entire z-stacks (Fig. S3). We calculated OPD for each point as the difference between $OPL_s$ for that point and $OPL_{med}$. The thickness of the sample at the same points where we obtained the OPD was determined using the corresponding confocal image. The intensity threshold for calculating thickness was set at the specific value based on the difference between the maximum and minimum intensity of the z-axis profile at the selected points (also see Figs. S2C, S5A, and S5C). Then, we calculated the dry mass density ("density" for short) for each point of the cytoplasm using the following formula:

$RI_s = RI_{med} + OPD/t$,

where $RI_{med}$ is the refractive index of the surrounding medium.

$RI_{med}$ was measured as 1.3362 (McCoy's 5A/10% FBS) and 1.3379 (DMEM/20% FBS). We determined the density of the sample from its RI using our calibration curves (Fig. 1D(iii); RI = $1.333 + 1.65 \times 10^{-4} \times C$, where C is density). We then obtained the mean density of selected points as a cytoplasm density of a cell. Also see "Measurement of Refractive Index (RI) of standard solutions"; Figs. 1D, 2B, and S2A.

To identify the density of the nucleus, we calculated the OPD of the nucleus and cytoplasm, $OPD_{nu\text{-}cy}$, for all the combination pairs of the selected points at cytoplasm and nucleus. We then obtained the average $OPD_{nu\text{-}cy}$ at each specific point. We corrected cytoplasm thickness using the thickness of cytoplasm (at the points that do not cross the nucleus and the points that cross the nucleus) as shown in Fig. S2B. Subsequently, we calculated the density at each point of nucleus using the measured $OPD_{nu\text{-}cy}$, $RI_{cy}$, the thickness of nucleus at that point. Finally, we derived the mean density of selected points as a nucleoplasm density of a cell. The process for measuring the density of chromosomes followed the same methodology as for nucleoplasm (see Fig. S5B-C). All measurements were done using FIJI software.

**Hypertonic and hypotonic treatment.**

For hypertonic treatment, cells were incubated in a medium supplemented with 0.9 mL DMEM and 0.1 mL 10X PBS just before observation. Cells were observed by OI-DIC within 1 h. For hypotonic treatment, cells were incubated in a medium supplemented with 1 mL DMEM and 1 mL MilliQ water just before observation. Cells were observed by OI-DIC within 1 h. The RI of medium: 1.3379 (McCoy's 5A/10% FBS, hypertonic), 1.3349 (McCoy's 5A/10% FBS, hypotonic) and 1.3403 (DMEM/20% FBS, hypertonic).

**Chromatin/DNA compaction analysis.**

To quantify DNA density, we used the obtained confocal images. In a central z-section of a mitotic cell (determined by visual inspection based on the largest area of cytoplasm), the DNA channel was denoised using a Gaussian blur filter ($\sigma = 2$) and threshold set using the Otsu dark method in FIJI. The resulting binary mask was converted into a selection, and the DNA mean fluorescence within this ROI was measured. All data points were normalized to the mean of unperturbed control cells. For hypotonic treatment, only images taken within 10 min after treatment were used.

**Chromatin preparation and condensation/droplet formation assay, and the droplet imaging.**

Fresh chicken blood was obtained from the wing vein of Tosa-jidori. Preparation of chicken native chromatin was as described previously (12). Briefly, 2 mL of fresh chicken blood was lyzed with 20 mL of MLB (60 mM KCl, 15 mM NaCl, 15 mM HEPES-KOH (pH 7.3), 2 mM $MgCl_2$, 0.1% NP-40, and 1 mM phenyl-methylsulfonyl fluoride [PMSF]) for 10 min on ice. After centrifugation at 1200 × g at 4 °C for 5 min, the supernatant was removed and resuspended in 20 mL of MLB. This step was repeated four times before the samples were ready for chromatin purification. Chromatin purification was carried out as described by (13), with some modifications. The nuclei (equivalent to ~2 mg of DNA) in nuclei isolation buffer (10 mM Tris-HCl, pH 7.5, 1.5 mM $MgCl_2$, 1.0 mM $CaCl_2$, 0.25 M sucrose, 0.1 mM PMSF) were digested with 50 U of micrococcal nuclease (Worthington, Lakewood, NJ) at 30 °C for 2 min. The reaction was stopped by adding ethylene glycol tetraacetic acid (EGTA) to a final concentration of 2 mM. After being washed with nuclei isolation buffer, the nuclei were lyzed with lysis buffer (10 mM Tris-HCl (pH 8.0), 5 mM EDTA, 0.1 mM PMSF) on ice for 5 min. The lysate was dialyzed against dialysis buffer (10 mM HEPES-NaOH (pH 7.5), 0.1 mM EDTA, 0.1 mM PMSF) at 4 °C overnight using Slide-A-Lyzer (66380 Thermo Scientific). The dialyzed lysate was centrifuged at 20400 × g at 4 °C for 10 min. The supernatant was recovered and used as the purified chromatin fraction. The purity and integrity of the chromatin protein components were verified by 14% SDS–PAGE (Fig. S7A). To examine the average DNA length of the purified chromatin, DNA was isolated from the chromatin fraction and electrophoresed in a 0.7% agarose gel (Fig. S7B).

Samples of chicken chromatin (4 μg) were incubated with 10 mM HEPES-NaOH (pH 7.5), 0.8 mM $MgCl_2$, 100 mM KCl, and indicated concentration of crowder in 200 μL of a reaction mixture at room temperature for 30 min. The mixture (10 μL) was mounted on a glass slide with 2 μL of DAPI solution (82 μg/mL). The coverslips were sealed with nail polish. The crowders used were polyethylene glycol (PEG; P5413, Sigma-Aldrich), Dextran (180~210 kDa, 041-22612, Fujifilm), and bovine serum albumin (BSA; BAC65, Equitech-Bio, Inc), and first dissolved in a buffer containing 10 mM HEPES-NaOH (pH 7.5) and 0.1 mM EDTA as 30% (w/v) solution. ImageJ was used to measure the droplet diameter by converting droplet images to binary images with a threshold of 80-255. Droplet diameters were then measured using "Analyze Particles".

For the chromatin assays at 40 mg/mL PEG, 200 mg/mL BSA, or 100 mg/mL Dextran (Panel 4 in Figs. 6A, 6C, and S7C), the chromatin droplets were first formed at 20 mg/mL PEG, 100 mg/mL BSA, or 50 mg/mL Dextran, and then additional crowder was added.

For the droplet dilution assays (Fig. 6D), chromatin droplets were first formed at 20 mg/mL PEG, 100 mg/mL BSA, or 50 mg/mL Dextran. Then, 9 volumes of a dilution buffer containing 10 mM HEPES-NaOH (pH 7.5), 0.8 mM $MgCl_2$, 100 mM KCl with or without crowder were added, incubated for 3 h and mounted with DAPI as described above.

For the chromatin droplet imaging, optical sectioning images were recorded with a 200 nm step size using a DeltaVision microscope (Applied Precision) and deconvolved to remove out-of-focus information. Projected images from five sections were shown as described previously (14).

**Immunostaining of ribosomal P protein.**

HCT116 cells expressing SMC2-mAID-mClover were grown on the poly-l-lysine–coated (P1524-500MG, Sigma-Aldrich) coverslips (C018001, Matsunami) for 2 days. All following processes were performed at room temperature. The cells were fixed with 1.85% formaldehyde (064-00406, Wako) on coverslips for 15 min, permeabilized with 0.5% Triton X-100 (T-9284, Sigma-Aldrich) for 5 min. After washing twice with HMK [20 mM Hepes (pH 7.5) with 1 mM MgCl2 and 100 mM KCl] for 5 min, the cells were incubated with 10% normal goat serum (NGS; 143-06561, Wako) in HMK for 30 min. The cells were incubated with diluted primary antibodies in 1% NGS in HMK for 1 hour: mouse anti-ribosomal p protein (1:10000 dilution; a gift from Dr. Uchiumi at Niigata University)(15). After being washed four times with HMK, the cells were incubated with diluted secondary antibodies in 1% NGS in HMK for 1 hour: goat anti–mouse IgG Alexa Fluor 594

(1:1000; A11032, Thermo Fisher Scientific), followed by washing four times with HMK. Cells were then stained with 4’,6-diamidino-2- phenylindole (DAPI) (0.5 μg/ml) (10236276001, Roche) for 5 min, followed by PPDI [20 mM Hepes (pH 7.4), 1 mM $MgCl_2$, 100 mM KCl, 78% glycerol, and paraphenylene diamine (1 mg/ml) (695106-1G, Sigma-Aldrich)] mounting. Optical sectioning images were recorded with a 0.5 μm step size using FV3000 confocal microscope equipped with an oil immersion 60X objective lens (Olympus UPlanXApo 60X NA: 1.42).

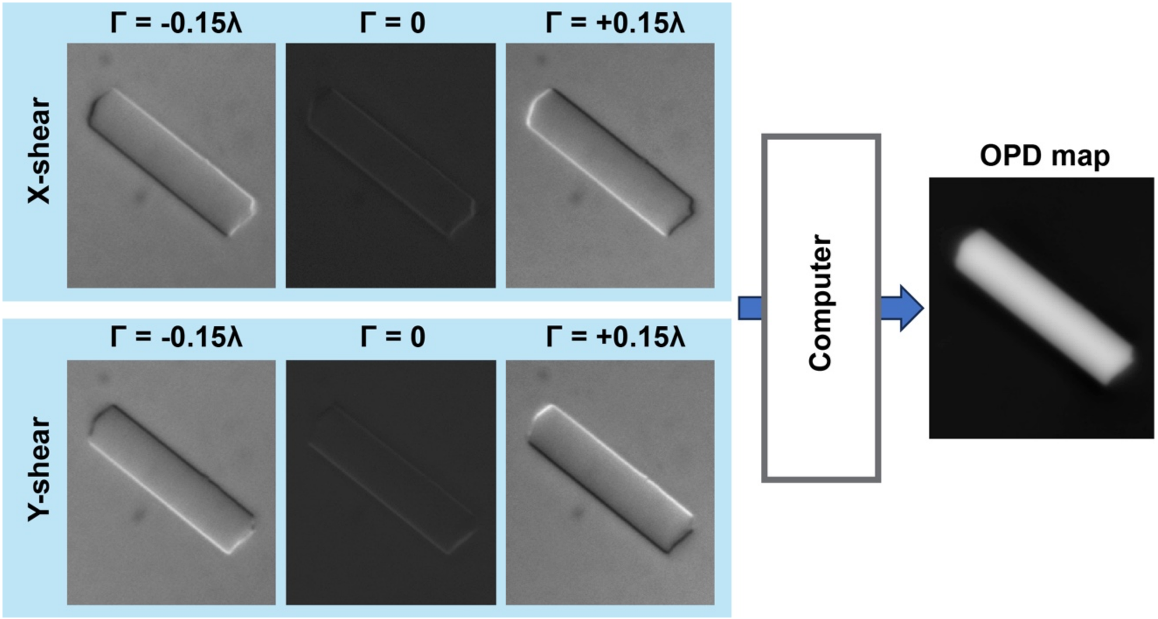


**Figure S1.**

**Principle of OI-DIC imaging**

OI-DIC can rapidly switch the shear directions of DIC without mechanically rotating the specimen or the prisms (8). A set of raw DIC images with orthogonal shear directions and different biases is captured within a second (Left). Specialized software computes a phase gradient vector map and then a quantitative phase image (Right).

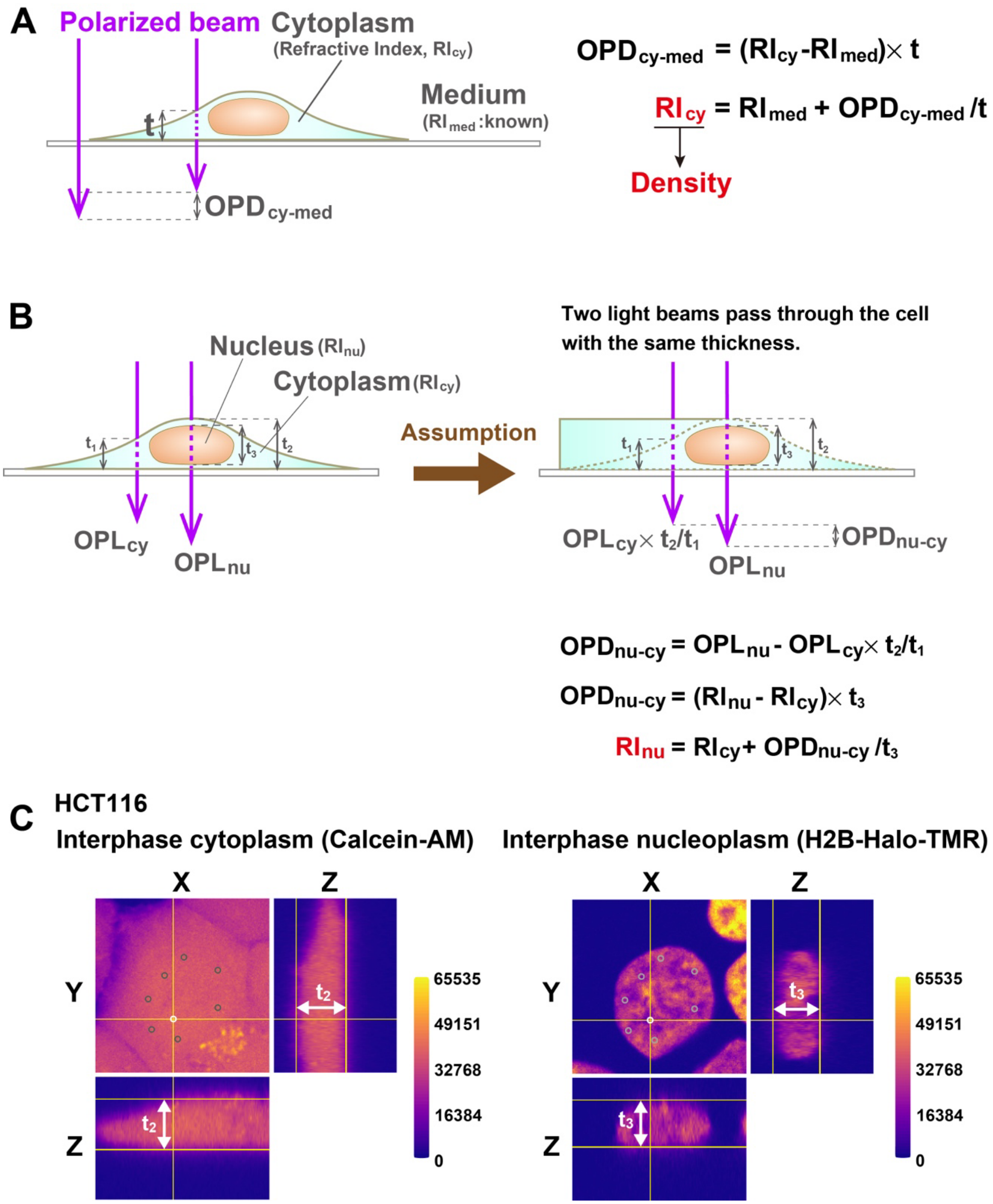


**Figure S2.**

**A simple schematic of the method used to estimate the refractive index (RI) in interphase cells.**

**(A)** The cytoplasm $RI_{cy}$ can be calculated based on the OPD measured by OI-DIC, the RI of medium ($RI_{med}$) measured by refractometer, and thickness (t) values. For details, see SI Materials and Methods. **(B)** A calculation scheme for the RI of the nucleus ($RI_{nu}$). We obtained $RI_{cy}$ based on the measured values for $OPD_{cy\text{-}med}$, thickness $t_1$, and $RI_{med}$. Because the thicknesses of the sample through each of the two polarized beams passed should be the same, we multiplied $OPL_{cy}$ by $t_2/t_1$ to obtain $OPD_{nuc\text{-}cy}$ ($t_2$, thickest point of the cell including nucleus and cytoplasm; $t_1$, thickness of cytoplasm region beside nucleus). $RI_{nu}$ was calculated based on $RI_{cy}$ and nuclear thickness $t_3$. **(C)** Thickness measurement of HCT116 cytoplasm (left) and nucleus (right) to estimate the molecular density of the nucleus. Several points in the nucleus were selected. The threshold for cytoplasm thickness was set at 33% or more for the difference between the maximum and minimum intensity of the z-axis profile of the selected points. For the nucleus, a threshold was also set at 33%. Here, points of thickness indicated by a white circle for the cytoplasm ($t_2$) or nucleus ($t_3$) are shown in orthogonal view. The LUT (Lookup table) is mpl-plasma. Also see (B).

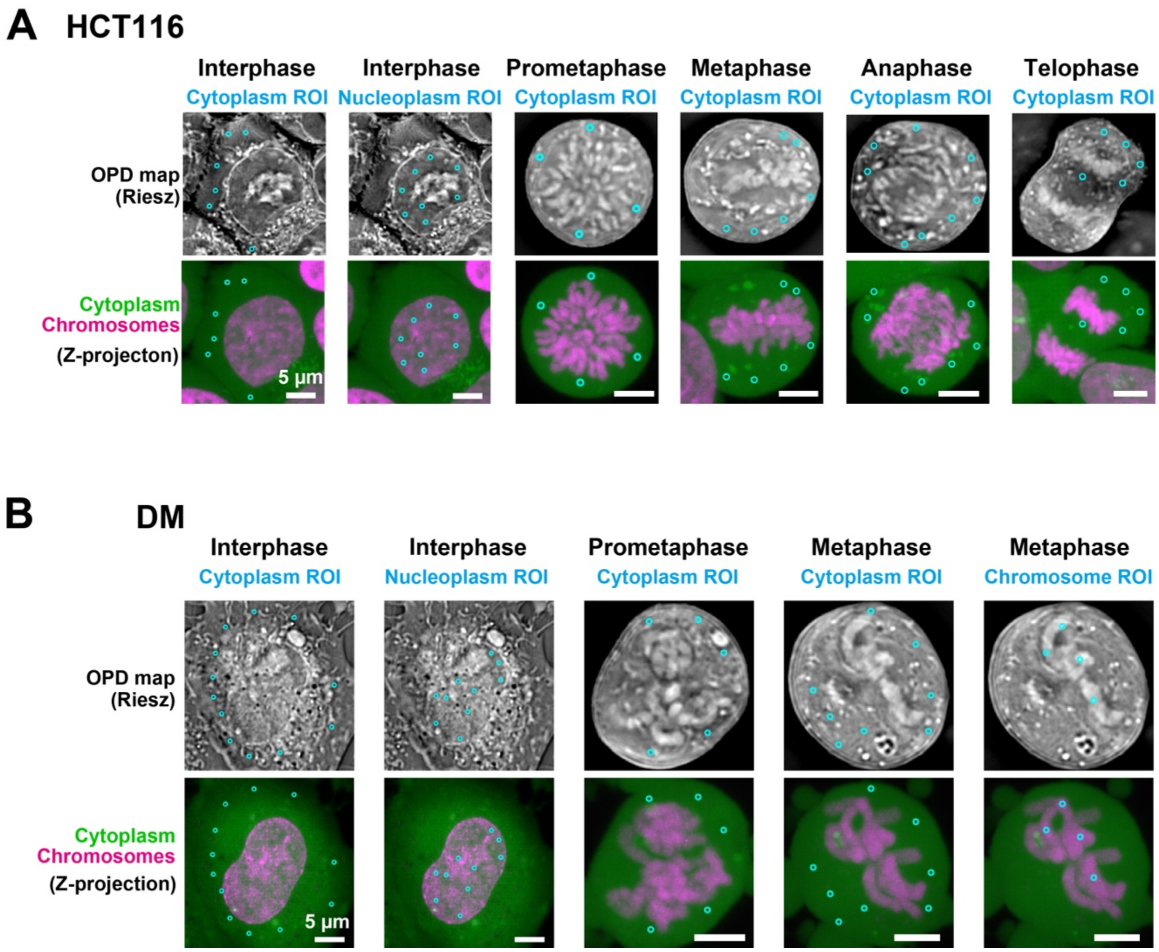


**Figure S3.**

**ROIs set to estimate total density.**

**(A)** Upper: The ROI set up to estimate total density overlaps the Riesz image of HCT116 cells. Note that the ROIs avoid apparent structures, presumably endoplasmic reticulum, Golgi apparatus, and mitochondria. Lower: The ROI set up to estimate total density overlaps the Z-projection of an entire cell. Note that the ROI avoids chromosomes in mitosis. **(B)** Upper: The ROI set up to estimate total density overlaps the Riesz image of DM cells. Note that the ROIs avoid apparent structures, presumably endoplasmic reticulum, Golgi apparatus, and mitochondria. Lower: The ROI set up to estimate total density overlaps the Z-projection of an entire cell.

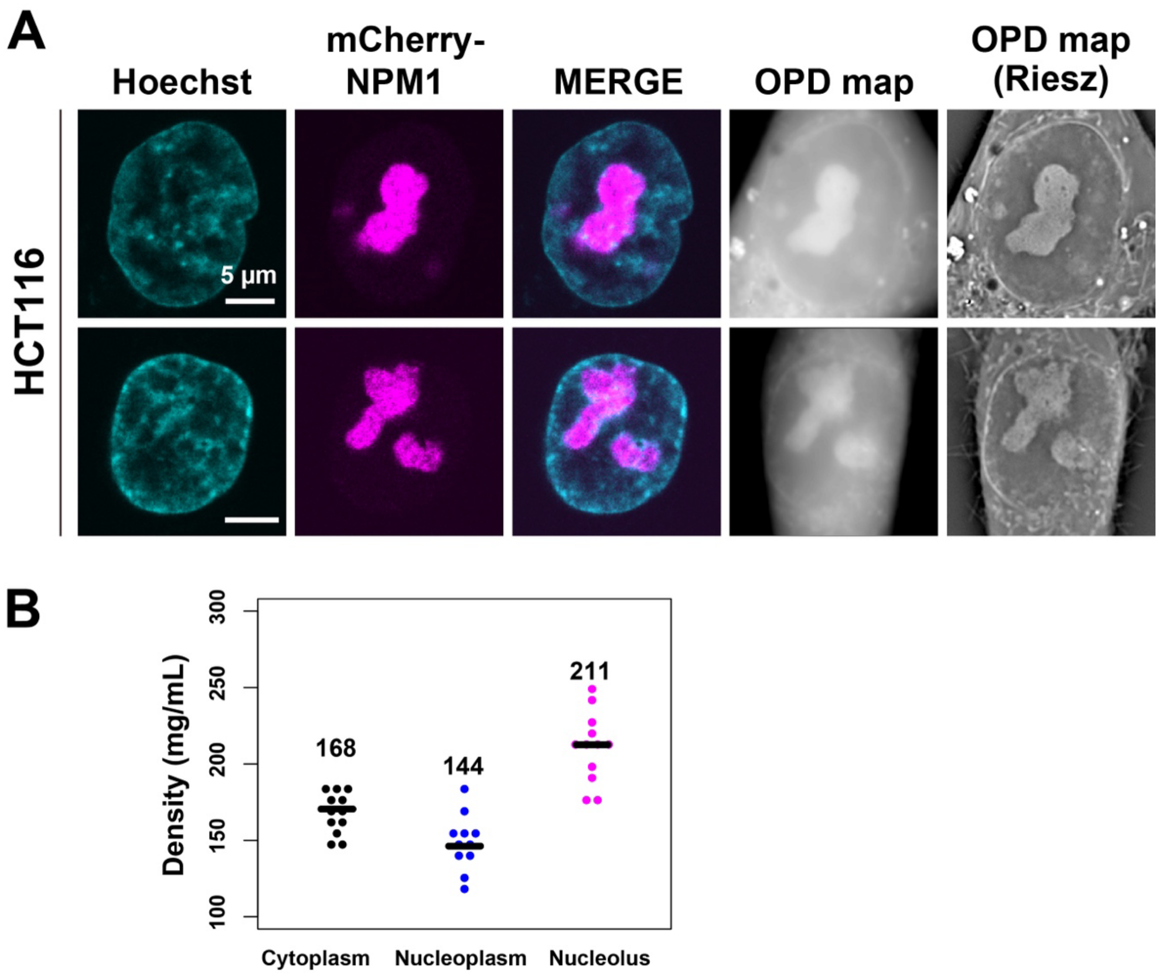


**Figure S4.**

**Density estimation in the nucleolus.**

**(A)** Confocal images of DNA nucleoli and OPD map and Riesz images obtained from OI-DIC imaging of live HCT116 cells. Riesz images are edge-enhanced OPD maps according to the inverse Riesz transform. mCherry-NPM1 was expressed in HCT116 cells. Scale bar: 5 µm. **(B)** Total molecular densities of interphase cytoplasm, nucleoplasm, and nucleolus in live HCT116 cells. Each dot represents the average of the estimated total density at several points within a single cell. For details, see Material and Methods. Black bars show the mean. Cell numbers are N = 12 (Interphase cytoplasm), N = 11 (Interphase nucleoplasm), and N = 12 (Nucleolus).

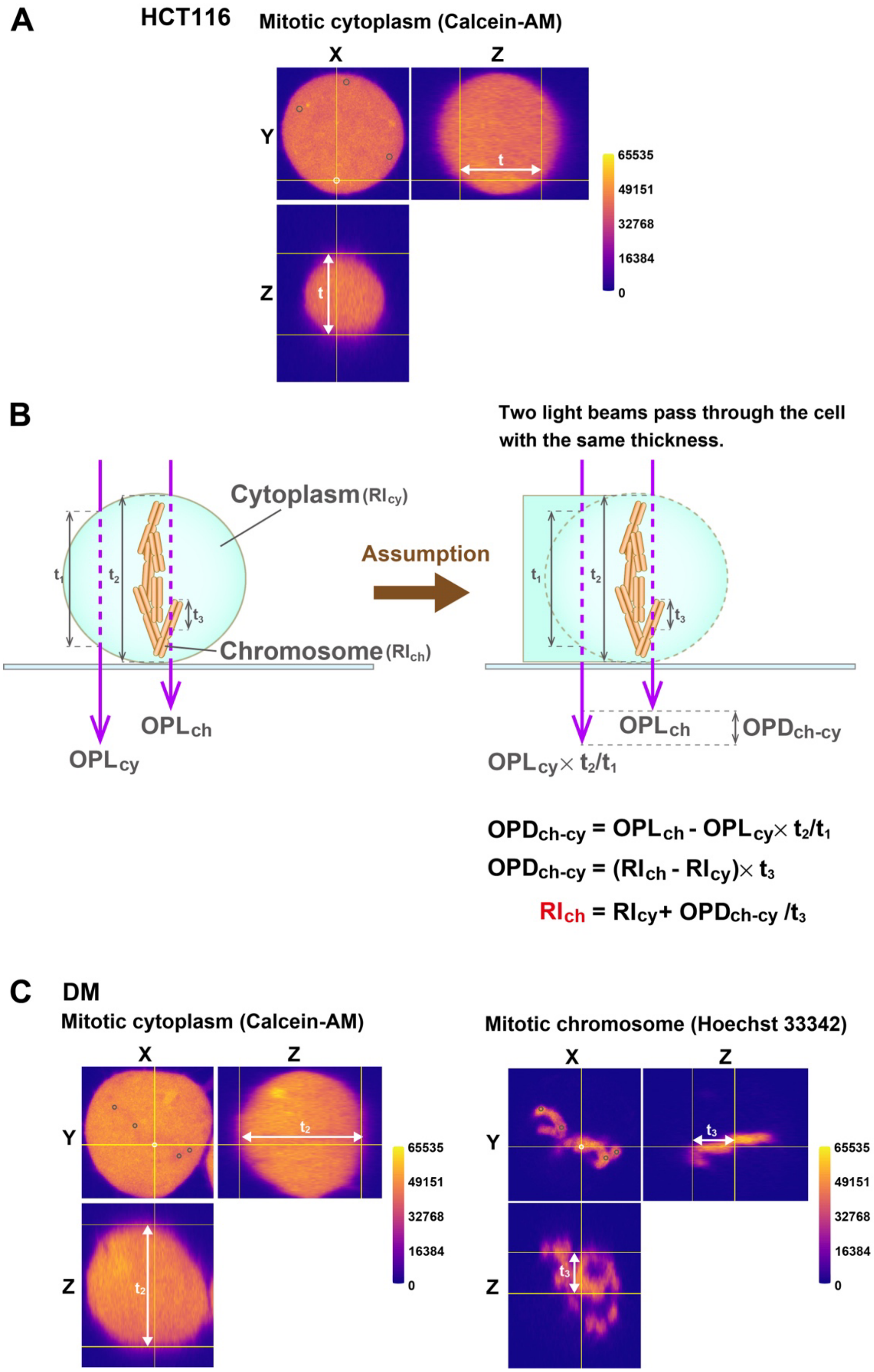


**Figure S5.**

**Simplified schematic of the method to estimate the RI in mitotic cells.**

**(A)** Thickness measurement of HCT116 cytoplasm. Several points that did not overlap with chromosomes were selected. The threshold for thickness was set at 40% or more of the difference between the maximum and minimum intensity of the z-axis profile of the selected points. Here, the thickness (t) of the point indicated by a white circle is shown in an orthogonal view. The LUT is mpl-plasma. Also see Fig. 2B. **(B)** A calculation scheme for the RI of chromosome ($RI_{ch}$). We obtained $RI_{cy}$ based on the measured values for $OPD_{cy\text{-}med}$, thickness $t_1$, and $RI_{med}$. Because the thicknesses of the sample through each of the two polarized beams passed should be the same, we multiplied $OPL_{cy}$ by $t_2/t_1$ to obtain $OPD_{ch\text{-}cy}$ ($t_2$, thickest point of the cell including chromosome and cytoplasm; $t_1$, thickness of cytoplasm region beside chromosome). $RI_{ch}$ was calculated based on $RI_{cy}$ and chromosome thickness $t_3$. **(C)** Thickness measurement of Indian Muntjac DM cytoplasm (left) and chromosomes (right) to estimate the molecular density of chromosomes. Several points were selected where chromosomes were present but not overlapping. The thickness threshold for cytoplasm was set at 40% or more of the difference between the maximum and minimum intensity of the z-axis profile of the selected points. For chromosomes, a threshold was set at 33%. Here, cytoplasm thickness ($t_2$) and chromosome thickness ($t_3$) of the point indicated by a white circle are shown in an orthogonal view. The LUT is mpl-plasma. Also see (B).

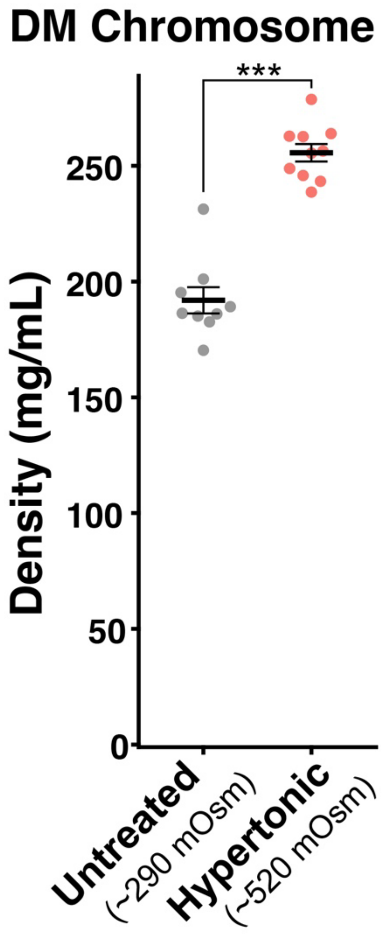


| DM cells | Density (mg/ml) |
|---|---|
| Untreated chromosome | 192 ± 17.0 |
| Hypertonic chromosome | 256 ± 17.0 |

**Figure S6.**
**Density imaging of chromosomes in live mitotic Indian Muntjac DM cells.**
Hypertonic treatment increases the total densities of chromosomes in live mitotic DM cells. Each dot represents the average of the estimated total density at several points of chromosomes within a single cell. For details, see Fig. S5B-C, and SI Materials and Methods. Black bars show the mean, error bars represent SE and cell numbers are N = 9 (Untreated) and N = 10 (Hypertonic). ***, $P < 0.0001$ by the two-sided unpaired t-test ($P = 1.9 \times 10^{-7}$).

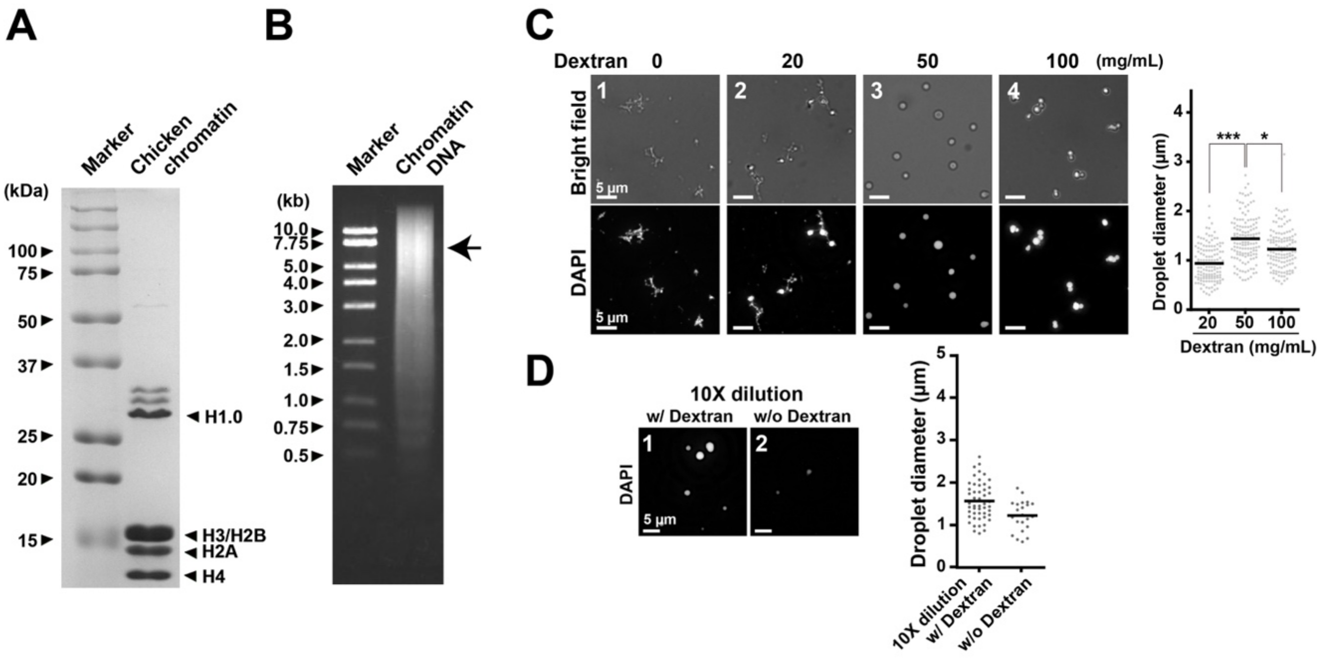


**Figure S7.**

**Chicken native chromatin preparation and droplet formation with Dextran.**

**(A)** SDS-PAGE of purified chicken native chromatin. Coomassie Brilliant Blue (R-250) staining shows that nucleosome core histones H2A/H2B, H3, and H4, as well as linker histone H1.0, are present in the sample. **(B)** Agarose gel of purified chromatin DNA. Average DNA length is ~6 kb (shown by arrow). **(C)** Formation of chromatin condensates/droplets with physiological salt and Dextran (~200 kDa). Fibrous chromatin condensates were induced by 100 mM $K^+$ and 0.8 mM $Mg^{2+}$ (Panel 1, 0 mg/mL Dextran). An increase in Dextran converted the fibrous condensates into droplets (Panels 2 and 3). A further increase in Dextran put the droplets together without fusions, suggesting the solid-like properties of the droplets (Panel 4). Diameters of formed droplets are shown on the right plots. ***, $P < 0.0001$ by Wilcoxon rank sum test for 20 mg/mL vs. 50 mg/mL PEG ($P = 7.2 \times 10^{-17}$). *, $P < 0.05$ for 50 mg/mL vs. 100 mg/mL PEG ($P = 2.0 \times 10^{-4}$). **(D)** The droplet dilution assay. After the droplets were formed with 50 mg/mL of Dextran, the reaction mixtures were diluted 10-fold in a buffer with (Panel 1) or without (Panel 2) 50 mg/mL of Dextran. Panel 2 depicts fewer and smaller droplets than Panel 1, suggesting that the droplets were dissolved in the diluted buffer without Dextran. A quantitative analysis of the droplet dilution assay (right). The amount and diameters of droplets from a randomly picked area ($6.0 \times 10^4$ µm$^2$) are plotted.

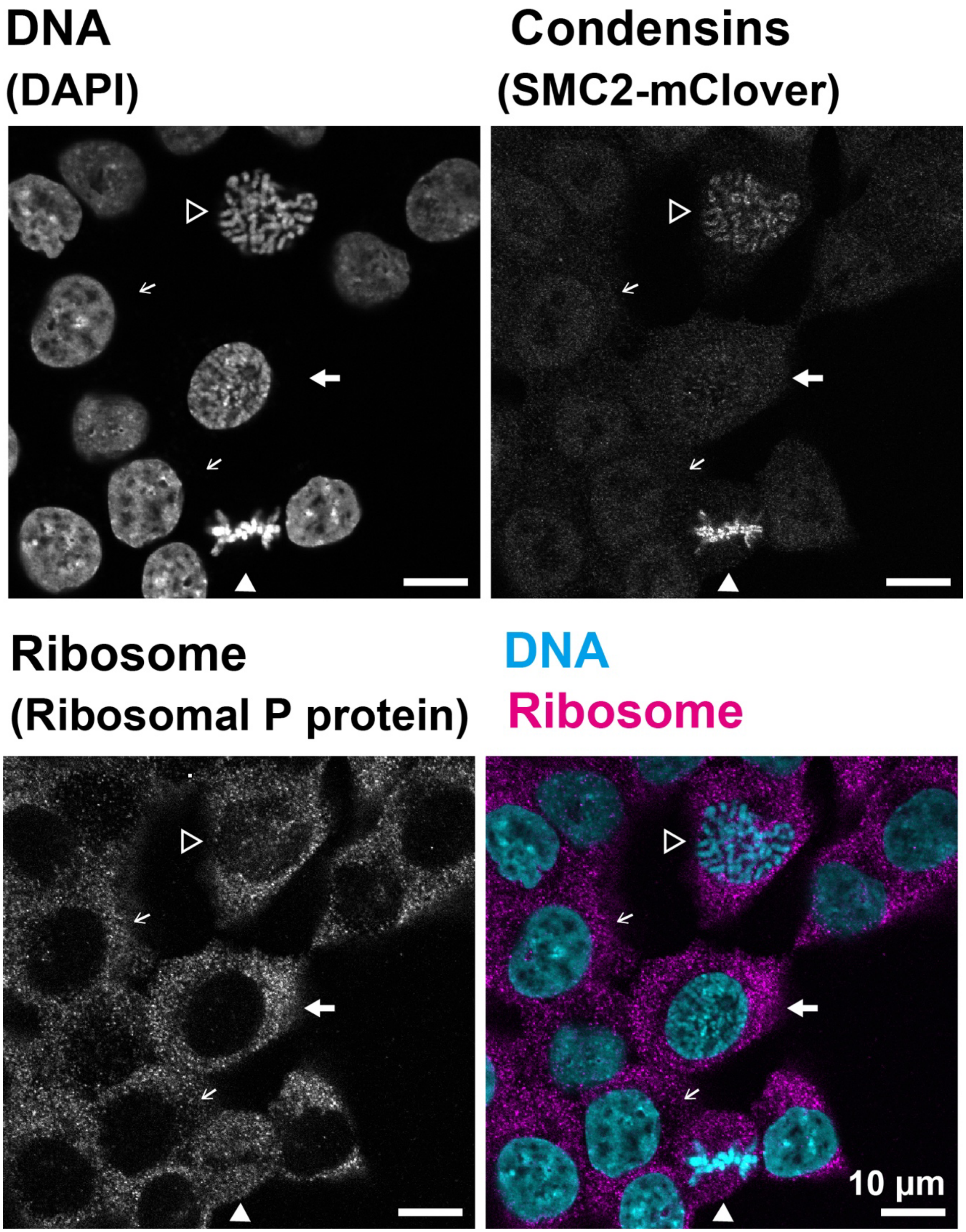


**Figure S8.**

**Localization of ribosomes by immunostaining.**

Confocal images of asynchronous HCT116 cells are shown. Interphase cells are indicated by thin arrows, prophase by a thick arrow, prometaphase by an outlined arrowhead, and metaphase by a filled arrowhead. Condensins are visualized by mClover-tagged endogenous SMC2, the common subunit of Condensins I and II. Note that during interphase and prophase, prior to nuclear envelope break down, ribosomes are sequestered from chromatin by the nuclear membrane. After nuclear envelope break down (prometaphase and metaphase), ribosomes can come into contact with chromosomes.

**Table S1.**
**Refractive index (RI) of HCT116 cells.**

| HCT116 cells | Refractive index (RI) |
|---|---|
| Interphase cytoplasm | 1.363 ± 0.006052 |
| Interphase nucleus | 1.356 ± 0.003834 |
| Prometaphase cytoplasm | 1.359 ± 0.002841 |
| Metaphase cytoplasm | 1.360 ± 0.002323 |
| Anaphase cytoplasm | 1.363 ± 0.003487 |
| Telophase cytoplasm | 1.358 ± 0.002984 |
| Mitotic cytoplasm (with hypertonic treatment) | 1.373 ± 0.003120 |
| Mitotic cytoplasm (with hypotonic treatment) | 1.348 ± 0.003112 |

**Table S2.**
**Refractive index (RI) of DM cells.**

| DM cells | Refractive index (RI) |
|---|---|
| Interphase cytoplasm | 1.354 ± 0.002060 |
| Interphase nucleus | 1.356 ± 0.002264 |
| Prometaphase cytoplasm | 1.360 ± 0.001850 |
| Metaphase cytoplasm | 1.361 ± 0.001938 |
| Mitotic cytoplasm (with hypertonic treatment) | 1.378 ± 0.002636 |
| Mitotic chromosomes | 1.365 ± 0.002810 |
| Mitotic chromosomes (with hypertonic treatment) | 1.376 ± 0.002238 |

**Movie S1 (separate file).**

Bright field movie of fusions of chromatin liquid droplets formed by 20 mg/mL of PEG (10 sec/ frame). Three fusion events were recorded in total.
